# Reproducible autosomal gene expression changes with loss of typical X and Y complement across tumor types

**DOI:** 10.1101/2025.09.24.678401

**Authors:** Seema B. Plaisier, Robert Phavong, Mason Farmwald, Mariah Lee, Teagen Allen, Malli Swamy, Ilsa Rodriguez, MacKenzie Wells, Nadia Phaneuf, Susan Christine Massey, Jared Del Rosario, Juvelyn Hart, Alexander Mangelsdorf, Martin Van Der Jagt, Alex R. DeCasien, Kenneth H. Buetow, Melissa A. Wilson

## Abstract

Although there are known sex differences in cancer incidence, severity, and treatment, the sex chromosomes are typically excluded from genomic analyses because of the unique technical challenges associated with assessing their copy number, sequence variation, and expression. Here we assess sex chromosome complement in three widely-used human genomics datasets from normal (non-cancerous) tissues, primary tumors, and cancer cell lines and study the effects on genome-wide gene expression. Expected sex chromosome complements based on reported patient sex were observed in non-cancerous tissues, but about half of tumors and cancer cell lines showed loss of typical sex chromosome gene expression across tissue types with three categories: loss of chromosome Y (LOY), loss of chromosome X (LOX) and reactivation of the inactive X chromosome (XaXa). Genes consistently differentially expressed in tumors with loss of chromosome X, loss of chromosome Y, or loss of X chromosome inactivation are associated with the hallmarks of cancer and include both sex-linked and autosomal genes from nearly all chromosomes, druggable genes, and genes with molecular functions relevant to cancer signaling, such as kinase activity. Strikingly, tumors that are X0, including tumors from female patients that have lost an X chromosome and tumors from male patients that have lost a Y chromosome, cluster together by gene expression profile. Patients with tumors that have LOX or LOY had poorer survival outcomes compared to those with tumors that had maintained their sex chromosome complement. Further, LOX and LOY eliminates nearly all of the differential gene expression between tumors from different patient sexes, affecting sex chromosomal and autosomal gene expression. Going forward, considering patient sex as well as the entire genome, including assessment of the sex chromosome complement, will provide additional insights into personalized tumor etiology, progression, treatment, and patient outcome.

**Teaser:** Loss of typical sex chromosome complement is present in primary tumors and cancer cell lines across tissue types. We identify consistent autosomal gene expression changes across multiple cancers when the sex chromosome complement is altered, and show that loss of sex chromosomes reduces tumor gene expression differences between patient sexes. We show that loss of sex chromosomes is also associated with poorer patient survival.

## Introduction

Sex differences in cancer have been observed in incidence, severity, progression, survival, and sensitivity to therapeutic interventions (Lopes-Ramos, Quackenbush, and DeMeo 2020; Rubin 2022; Haupt et al. 2021). The presence of different sex chromosome complements (typically XX or XY) are a key genetic difference in tissues from karyotypical females and males, with all individuals possessing at least one X chromosome. Given that the sex chromosomes contain hundreds of genes that are involved in cellular processes that can affect cancer progression (Spatz, Borg, and Feunteun 2004), such as immunity and tumor suppression, and that sex chromosomes regulate the expression of autosomal genes (Balaton et al. 2018; Lopes-Ramos, Quackenbush, and DeMeo 2020; San Roman et al. 2024), the sex chromosome complement is very likely to contribute to sex differences in disease onset, progression, and treatment. Regulatory mechanisms specific to sex chromosomes, such as X chromosome inactivation, may influence cancer incidence; for example, tumor suppressor genes on the X chromosome that escape X chromosome inactivation are thought to contribute to the observed reduced cancer rate in XX females (Dunford et al. 2017). Despite this, many studies do not assess or analyze the sex chromosome complement (SCC) of the samples they are analyzing (Wilson and Buetow 2020; Wise, Gyi, and Manolio 2013), effectively excluding 5% of the genome, and the most common genetic variation in human populations.

Assessing the sex chromosome complement in a sample provides information that is relevant to disease risk and severity. Sex chromosomes can be lost from individual cells during the normal aging process (Gadek et al. 2025; Guttenbach et al. 1995; Jacobs et al. 1963; Russell et al. 2007) and such loss is linked to a worse prognosis in some cancers (Abdel-Hafiz et al. 2023; Cáceres et al. 2020; Kido and Lau 2015; Brown and Machiela 2020; Forsberg et al. 2014). In normal aging, both the X and the Y chromosomes experience an age-dependent loss in lymphocytes, with up to 1.34% of males ages 76 to 80 exhibiting chromosome Y loss, and roughly 5% of females ages 75 and above exhibiting loss of chromosome X (Guttenbach et al. 1995). Frequency of loss of chromosome X from routine cytogenetic analysis was shown to range from 0.07% in females under 16 years of age to 7.3% in females over 65 years of age (Russell et al. 2007). Hematopoietic mosaic loss of chromosome Y has been associated with an increased risk for cardiac pathology (Sano et al. 2022). Y chromosome loss is associated with genome instability, higher tumor mutational burden, differential expression of tumor suppressors and oncogenes, and lower survival in several cancer types (Müller et al. 2023; Qi et al. 2023). Bladder cancer cells that lost chromosome Y were more aggressive as they were able to evade the immune system by altering T cell functions (Abdel-Hafiz et al. 2023). Loss of chromosome X has also been linked to cancer phenotypes. Loss of whole chromosome X was predictive of lower event-free survival in pediatric neuroblastoma (Parodi et al. 2019). As sex chromosomes can be lost in normal aging or disease states, the reported sex of an individual may not indicate what sex chromosomes are present in patient tissues.

Expression of genes on the sex chromosomes can be used to detect the presence of transcriptionally active sex chromosomes, and a lack of expression can be used to infer sex chromosome loss. The X chromosome contains hundreds of genes that are essential for growth and development of cells, and as such, having at least one copy is necessary for proper cell growth and development (T. Wang et al. 2015). The X chromosome carries roughly 1500 genes (Spatz, Borg, and Feunteun 2004), of which 841 are protein-coding (Dyer et al. 2025). The long, non-coding transcript *XIST* is responsible for inactivation of all but one copy of the X chromosomes in cells where two or more X chromosomes are present. When two X chromosomes are present in a cell, transcription of *XIST* will dramatically increase. The *XIST* transcripts coat all but one X chromosome; the X chromosomes coated in *XIST* transcripts will get tightly compacted, repressing most gene expression from the inactivated X chromosomes (Cerase et al. 2015; Penny et al. 1996; Balaton et al. 2018; Chang et al. 2006; Lyon 1999; Lin et al. 2025; Cheng and Disteche 2004; Tukiainen et al. 2017). While X chromosome inactivation is expected in all cells with more than one X chromosome, loss of this dosage compensation mechanism has been observed in induced pluripotent stem cells from female donors in which low levels of *XIST* correlated with increased protein expression of X-linked genes (Brenes et al. 2021). The Y chromosome has 693 genes (about 106 protein-coding), including many that are homologs of genes on the X chromosome (Rhie et al. 2023; DeCasien et al. 2025; Blanton et al. 2024). Y chromosome genes that are involved in sex determination early in development have been well characterized, but chromosome Y genes have been shown to be expressed across all adult tissues with functions outside of gonad differentiation (Godfrey et al. 2020; Maan et al. 2017). Although expression of these sex chromosome genes has been used to predict sex for specific applications (Teixeira et al. 2019), sex chromosome inference is not routinely performed in genomics analysis despite the important roles that sex chromosomes play in cell biology.

Here we develop and apply a method to use the expression of genes on the sex chromosomes to infer the functional sex chromosome complement of samples from commonly used human genomics datasets and show how differences in sex chromosome complement affect the global gene expression profile. To investigate the functional sex chromosome complement of tissue samples in the context of the reported sex of each individual, this study examines the expression of genes on the X and Y chromosomes in three large-scale publicly available data sets: (1) normal adult tissues from 54 organs from 946 adult individuals without any known disease diagnoses from the Genotype-Tissue Expression (GTEx) consortium (GTEx Consortium 2013; Carithers and Moore 2015; GTEx Consortium 2020); (2) over 20,000 primary tumor tissues and matched adjacent normal tissues spanning 33 cancer types from The Cancer Genome Atlas (TCGA) (Cancer Genome Atlas Research Network et al. 2013); and (3) 1019 cancer cell lines routinely used in controlled, preclinical studies from the Cancer Cell Line Encyclopedia (CCLE) (Ghandi et al. 2019; Barretina et al. 2012; Tsherniak et al. 2017). We find that functional sex chromosome complement generally matches the typical karyotype for the reported sex of the individual in healthy GTEx tissues, but between up to 56% of samples exhibit loss of functional sex chromosome expression in tumors (from TCGA) and cancer cell lines (from CCLE). We find that, when grouping according to differences in reported patient sex and observed sex chromosome complement, autosomal genes known to be involved in the cancer are consistently differentially expressed across tumor types, suggesting a regulatory effect of each sex chromosome affecting the entire genome. We observe that sex differences in gene expression are greatly reduced in tumors that have a single X – via loss of X or loss of Y – highlighting the contribution of genetics in driving sex differences in gene expression. Finally, we confirm lower survival for XY males compared to XX females, but find, surprisingly, that the patient sex difference in survival disappears with the loss of a sex chromosome with all of those patients showing equally poor survival.

## Results

### Transcription based approach to infer the functional sex chromosome complement

Our approach to study sex chromosomes in tissues with and without cancer begins with the expression of sex chromosome genes that can indicate which sex chromosomes are present and functionally (transcriptionally) active. We identify a set of genes on chromosomes X and Y that we verified are expressed in normal adult tissues (Figure 1 and Methods). The expression of *XIST* was used to infer the presence of at least two X chromosomes in which one is silenced (XaXi). Lack of *XIST* expression was interpreted as loss of an X chromosome when DNA ploidy shows a single copy of X (LOX) or as X chromosome reactivation when the DNA ploidy shows two copies of X (XaXa). The expression of two or more of the set of Y chromosome genes was used to infer the presence of an active Y chromosome. The expression of Y chromosome genes was generally correlated, but *RPS4Y1* had high expression throughout the samples from male patients, so we required expression of at least one additional chromosome Y gene to indicate the presence of an active Y chromosome (Figure 1, Figure S1). Using these genes together allowed the inference of the functional sex chromosome complement (SCC), which was used to study the differences between sample groups with typical and atypical SCCs given the reported sex of the individual. Using a TPM expression threshold allowed for a clear, reproducible definition of the expression of a gene in a particular sample.

**Fig. 1.**
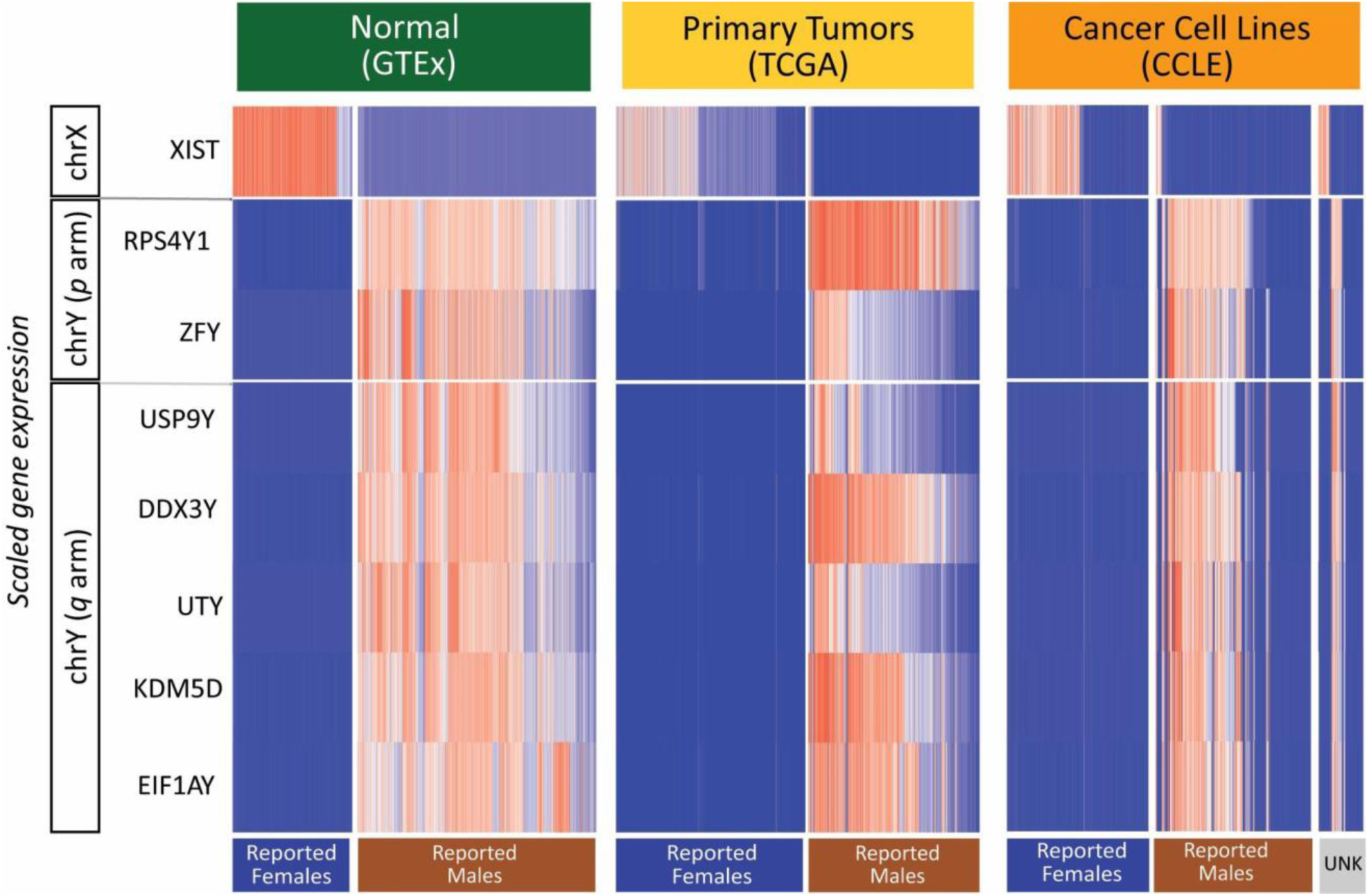
Sex chromosome gene expression for sex chromosome inference. Heatmaps showing expression of genes on the sex chromosomes used for inference of the sex chromosome complements in normal tissues (GTEX), primary tumor tissues (TCGA), and cancer cell lines (CCLE). Expression for each gene is scaled within each data set, with highest expression shown in red and lowest expression shown in blue. Samples are grouped by the reported sex of the person from whom the original sample was isolated (the patient from which the cell line was derived in the case of cancer cell lines).

### Sex chromosome gene expression across tissue types in the absence of cancer

This approach for inferring sex chromosome complement was applied to normal (without cancer) adult tissues (Figure 2, Figure S3) as a basis of reference for cancer tissues. In normal adult tissues (GTEx), the distribution of *XIST* expression was centered well above our threshold for high expression (>10 TPM) in the majority of tissues (Figure 2 A). We observe that 87% of all samples from subjects reported to be female had high expression of *XIST* (>10 TPM). We observed that the mean of the distribution of *XIST* expression fell below our threshold for expression (>10 TPM) in whole blood as well as all regions of the brain except the cerebellar hemisphere and cerebellum. Of the 13% of GTEx samples that showed loss of *XIST* expression, 48% were from non-cerebellum brain regions (361/750) and 31% were from whole blood (232/750). Tissue types where the distribution of *XIST* expression was lower than the mean for other tissues also exhibited lower than average expression of chromosome Y genes (*DDX3Y* expression is shown as an example in Figure 2 B). These patterns suggest that using the same thresholding approach may not be appropriate for these tissues in their normal (non-cancerous) state. After excluding tissues with low overall expression of sex chromosome genes, 96.5% of normal tissue samples from female subjects had *XIST* expression and no expression of chromosome Y genes, indicating a XX genotype, and 99% of normal tissue samples from male subjects had expression of chromosome Y genes and no expression of *XIST*, indicating an XY genotype (Figure 2). Observing the inferred SCC for each tissue, the majority of tissues showed above 95% expected SCC (XaXi in female patient samples and XY in male patient samples) (Figure S3). Using our thresholds for expression, sex chromosome alterations were observed in only 3.4% of female tissue samples and 0.007% of male tissue samples in the absence of cancer (Figure 2).

**Fig. 2.**
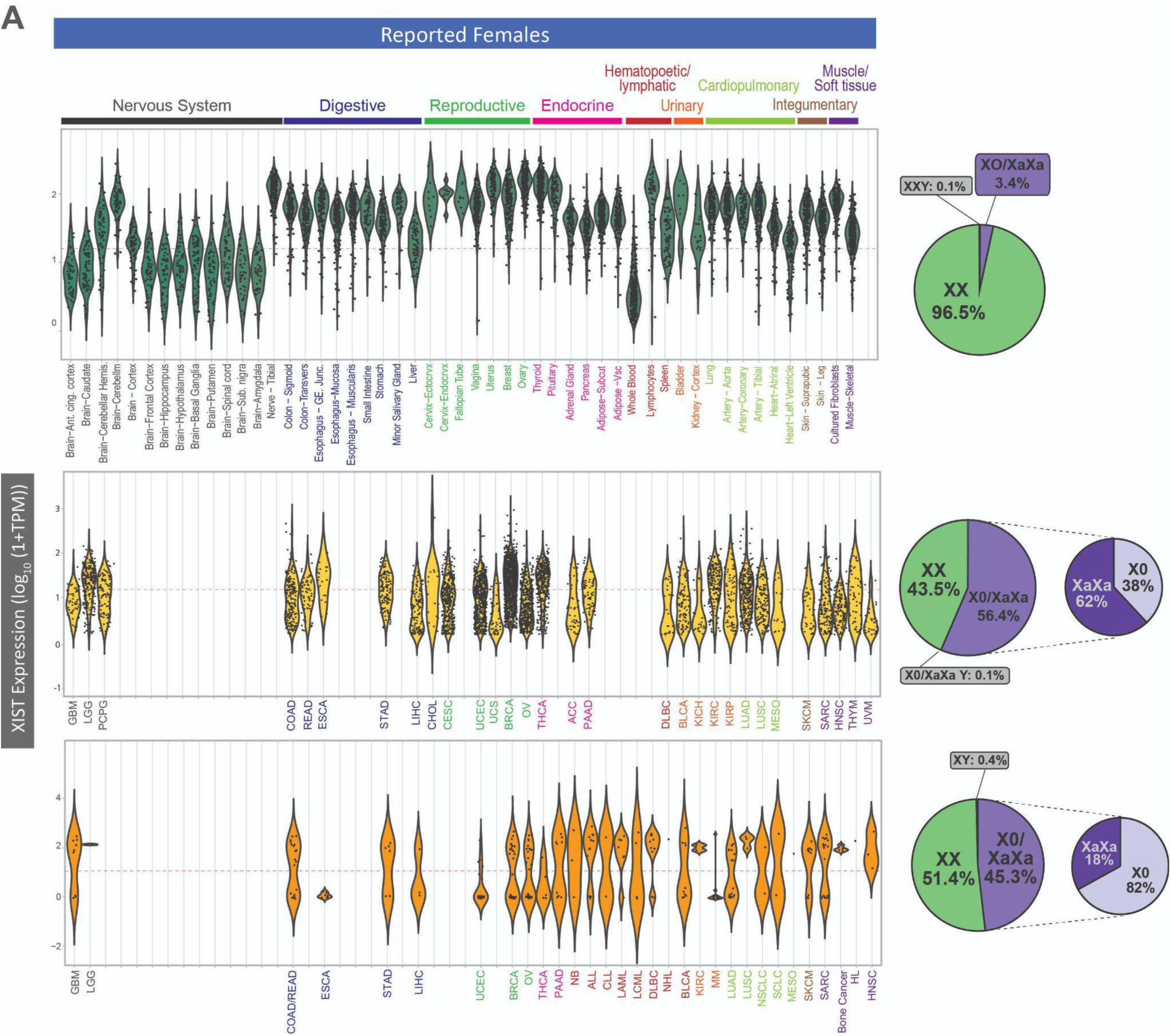
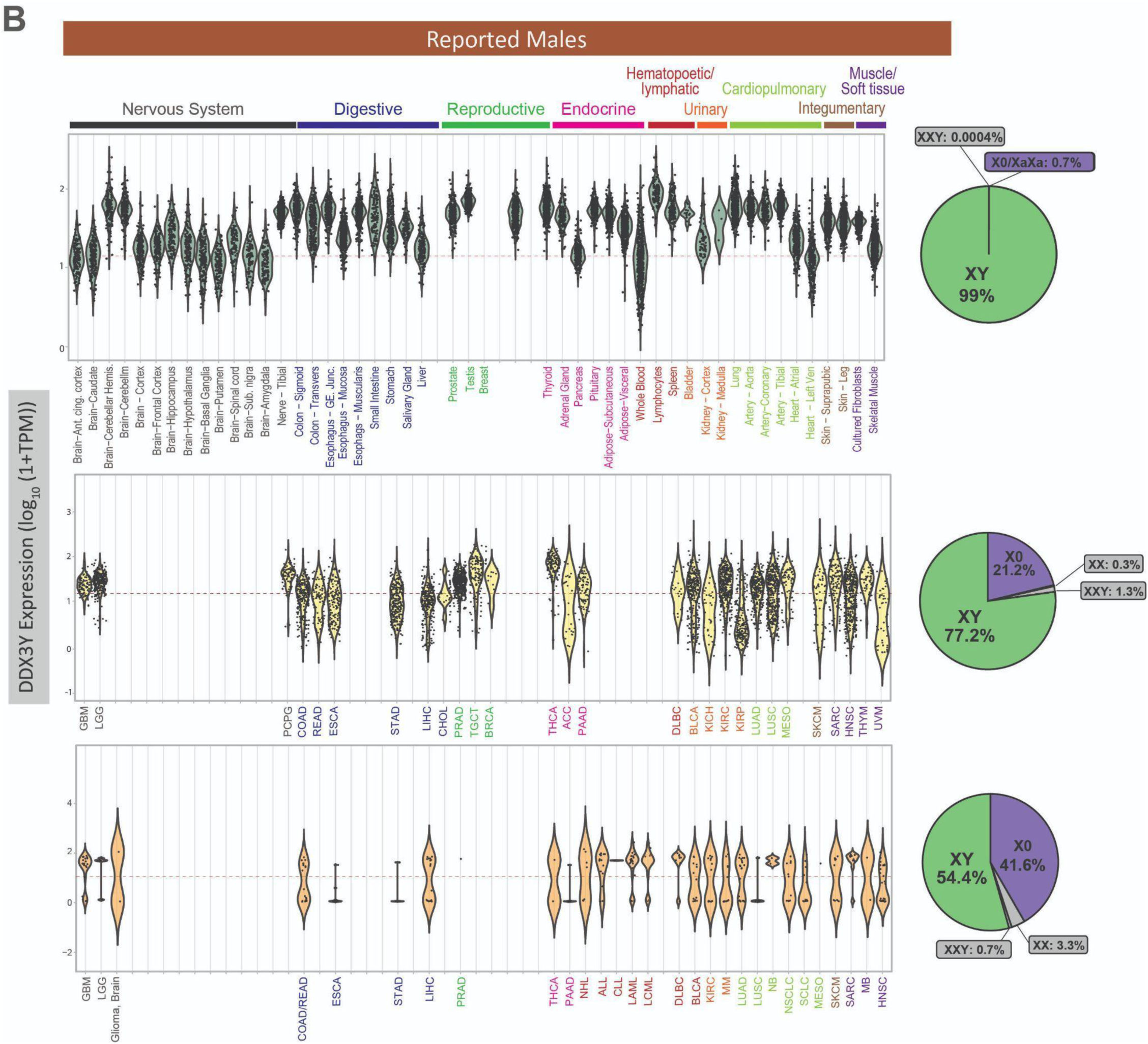
Sex chromosome loss in tumor tissues and cancer cell lines compared to adult normal tissues across many tissue types. (**A**), XIST expression and (**B**), DDX3Y expression across tissue types in GTEx (normal), TCGA (primary tumors) and CCLE (cancer cell lines). Red dashed lines show the minimum threshold for XIST or DDX3Y being called as expressed.

In addition to tissues from individuals without cancer (GTEx), we also examined marker gene expression in adjacent normal tissue collected from cancer patients (Figure S4). In adjacent normal tissue samples from female patients, 51% (188/370) of tissues expressed *XIST*, 42% (157/370) had low expression of *XIST*, and 7% (25/370) showed no expression of *XIST*. In adjacent normal tissue from male patients, the vast majority of tissues (96%, 340/353) expressed 2 or more chromosome Y marker genes, with only 3.6% (13/353) that showed did not show expression of at least two chromosome Y genes. When comparing pairs of normal adjacent tissue and tumor tissue from the same patient, *DDX3Y* tends to show lower expression in tumor tissues compared to paired adjacent normal tissue, showing an overall decrease in expression in many cancer types (Figure S4 B and C). By contrast, *XIST* expression is seen at higher, similar, and lower levels in tumors compared to paired adjacent normal tissue (Figure S4 A and C). This indicates that the majority of normal adjacent tissues from male patients had presence of an active Y chromosome and tumor tissues from the same patients lost chromosome Y if there was any change in the SCC. *XIST* loss is observed in varying degrees in normal adjacent tissues with no clear tendency towards loss or reactivation in tumor compared to adjacent tissue.

### Sex chromosome complements often do not match reported sex in tumor tissues and cancer cell lines

We observed more atypical sex chromosome gene expression in primary tumor tissues and cancer cell lines than observed in normal adult tissues (Figure 2, Figure S3). Over half of the primary tumor samples from female patients are inferred to have atypical X chromosome expression and about a quarter of samples from male patients have atypical Y chromosome expression (Figure 2). Among primary tumor samples from reported females with *XIST* expression below our expression threshold (10 TPM), 62% have XX DNA ploidy measurements from the same tumor, suggesting that both X chromosomes are active (XaXa), while the remaining 38% have single X DNA ploidy, consistent with loss of all but one X chromosome (X0) (Figure 2). Primary tumors show a wide distribution of expression of sex chromosome marker genes, which may reflect different amounts of sex chromosome loss in different cell types of tumors and/or a range of tumor purity in the TCGA samples. The proportion of atypical sex chromosome expression varies substantially across tumor tissue types (Figure S3). In tumors from female patients, uveal melanoma (UVM) has the lowest proportion of XaXi tumors from female patients at only 9% (3 out of 35), while thyroid carcinoma (THCA) has the highest proportion of XaXi tumors at 70% (260 out of 369). In tumors from male patients, kidney renal papillary cell carcinoma (KIRP) has the lowest proportion of typical XY at only 24% (51 out of 213) while glioblastoma (GBM) has one of the highest proportions of typical sex chromosome complement, at 99% XY (101 out of 102), next to breast cancer, with only 12 tumors from male patients in TCGA, which were all XY.

While primary tumors show heterogeneity for loss of typical X and Y expression, cancer cell lines generally exhibit bimodal distributions across tissue types (Figure 2). This bimodal distribution is also visible at the protein abundance level for chromosome Y genes measured by quantitative mass spectrometry (Figure S5 C, Figure S6). Similar measurements could not be conducted with *XIST*, because it is a long non-coding RNA that is not transcribed into a protein. When inferring sex chromosome complement, we infer that 45.3% of cancer cell lines derived from a tumor from a female patient have lost *XIST* expression. However, in contrast to primary tumors, the majority (82%) of cell lines exhibit single X ploidy indicating that one X chromosome was lost either in the tumor from which this cell line was derived or as the cell line was established and passaged, while the only the remaining 18% have XX ploidy indicating that both X chromosomes are active (XaXa) (Figure 2). In cancer cell lines from male patients, 41.6% of cell lines have lost the Y chromosome, which is nearly double the percentage of loss observed across primary tumors. For the majority of tissues of origin, CCLE contained cancer cell lines that had both expected and atypical sex chromosome complements (Figure S3). The proportion of cell lines that show evidence of sex chromosome loss in each tissue does not correlate to the proportion of tumors observed to have sex chromosome loss. For example, thyroid cancer primary tumors from female patients in TCGA showed the highest proportion of XaXi, but only 1 of the 5 thyroid cancer cell lines from female patients was inferred to be XaXi. As many genetic changes take place during the establishment of cancer cell lines, being aware of the sex chromosome complement may help to create a better matching cell line model to the cancer being studied.

For cancer cell lines in the CCLE, 88 of 671 cancer cell lines derived from primary tumors were not annotated with the sex of the patient from which the cell line was established. We inferred the sex chromosome complement of these unannotated cell lines and found that for 21 samples we infer an XX karyotype and for 25 we infer an XY karyotype, suggesting that we can predict the sex of the patient from which the cell line was derived. For the remaining samples, we do not detect *XIST* or chromosome Y gene expression, suggesting an X0 karyotype from loss of X from XX female patients or a loss of Y from XY male patients; we cannot infer whether the sample came from a patient with XX or XY chromosomes (Figure 1).

Within each of the data sets, we observed samples with sex chromosome gene expression patterns that may be representative of true biology or may be due to mislabeling, sample swapping, or contamination (Figure 1, Figure 2, Figure S3, Table 4). In GTEx normal adult tissues, three samples from reported females had gene expression that suggests an XXY genotype, as they have high expression of *XIST*, *RPS4Y1,* as well as *DDX3Y* or *KDM5D*, and low expression of the other chromosome Y genes. These samples are from three separate individuals and all other samples from the same individuals show XX expression patterns, so it is unlikely that these individuals have a germline sex chromosome aneuploidy. Similarly, four male GTEx samples have XXY expression patterns (high *XIST* and chromosome Y gene expression), occurring in only one sample from 4 separate individuals. In TCGA primary tumor samples, 16 samples from males have an inferred XX expression pattern (high *XIST* expression, chrY genes not expressed) and 66 have an inferred XXY expression pattern (high *XIST* expression, high chrY gene expression). However, male cancers with *XIST* expressed have been observed in a subset of tissue types, in some cases with molecular evidence of X chromosome inactivation such as reduced chromatin accessibility and increased DNA methylation (Sadagopan et al. 2022). Of the *XIST*-expressing male samples in TCGA, 77% (62 out of 82) were from testicular germ cell tumors. Of those, 12 were previously reported to have greater than two X chromosomes (Sadagopan et al. 2022) using DNA level measurements. In cancer cell lines from the CCLE, one cell line derived from a tumor from a female patient has an XY expression pattern and ten cell lines derived from a tumor from a male patient show XX expression pattern. Two cell lines from a male patient show an XXY expression pattern (HT1197_URINARY_TRACT, U266B1_HAEMATOPOIETIC_AND_LYMPHOID_TISSUE). Observations specific to tissue samples or cell lines with these unexpected sex chromosome complements should be analyzed with more scrutiny.

### Loss of sex chromosomes associated with increased age

Across all three datasets, samples inferred to have functional loss of a sex chromosome were slightly older on average compared to those who maintained their expected sex chromosome complement. Age was significantly correlated with both *XIST* expression and average Y chromosome marker gene expression (Figure S7). In normal, non-cancerous tissues (GTEx), the small number of female samples that had lost *XIST* expression showed a higher average age by 2.5 years (thought this was not statistically significant). Normal, non-cancerous male samples that had low chromosome Y gene expression (none had no chromosome Y gene expression) showed a higher average age by 5.1 years (Student’s t-test p-value < 0.0001). In primary tumor tissues from TCGA, female patients with tumors that had lost *XIST* expression were 3.7 years older on average (p-value < 0.0001) and male patients with tumors that had lost chromosome Y gene expression were 4.6 years older on average (p-value < 0.0001). For cancer cell lines that had lost *XIST* expression, female patients with the tumors those cell lines were generated from were 5.4 years older on average (not statistically significant). For cell lines that had lost chromosome Y gene expression, male patients with the tumors those cell lines were generated from were 12.2 years older on average (p-value = 0.0006). In all datasets, loss of *XIST* and Y gene expression occurred across the age range and was not limited to older individuals, but loss of *XIST* and chromosome Y gene expression was associated with a higher age (Figure S7).

### Genes differentially expressed based on sex chromosomes are associated with cancer functions

To understand global effects of changes in sex chromosome complement in human tumors, we conducted differential expression based on patient reported sex and sex chromosome complement and identified genes that were changing consistently across tissue types. As age had a significant influence on *XIST* and chromosome Y gene expression, age was included as a covariate in the differential gene expression analysis. For each tissue type, differential gene expression profiles were calculated comparing two sample groups as long as there were at least three samples in each group. For samples from female patients, samples were split into groups based on both single X ploidy and *XIST* expression. Gene expression profiles were calculated based on three sample groups within each tissue type: (1) XX ploidy at the DNA level and *XIST* gene expression which indicates the typical XX genotype where one X is inactivated (female, XaXi), (2) XX ploidy with no *XIST* expression indicating XX genotype with two active X chromosomes (female, XaXa), and (3) single X ploidy and no *XIST* expression indicating loss of all but one X chromosome (female, X0). Each of the two atypical sex chromosome complements was compared to samples with the typical XaXi karyotype. For samples from male patients, a loss of Y (LOY) gene expression profile was calculated between samples that had no expression of chromosome Y genes and no expression of *XIST* (male, X0) compared to those that had expression of chromosome Y genes and no expression of *XIST* (male, XY). We also assessed genes that are consistently differentially expressed between XaXi female patient tumors and XY male patient tumors. In all four cases, we constructed consistent gene signatures with genes that were significantly differentially expressed (p-value < 0.05) in at least 60% of the tissues and were changing in the same direction in all tissues (Figure 3). While we filtered using statistically significance and direction to identify consistent gene changes that had a higher likelihood of affecting tumor cell biology, the full gene expression profiles of sex chromosome loss or X reactivation were generally consistent across tissues (Figure S8).

**Fig. 3.**
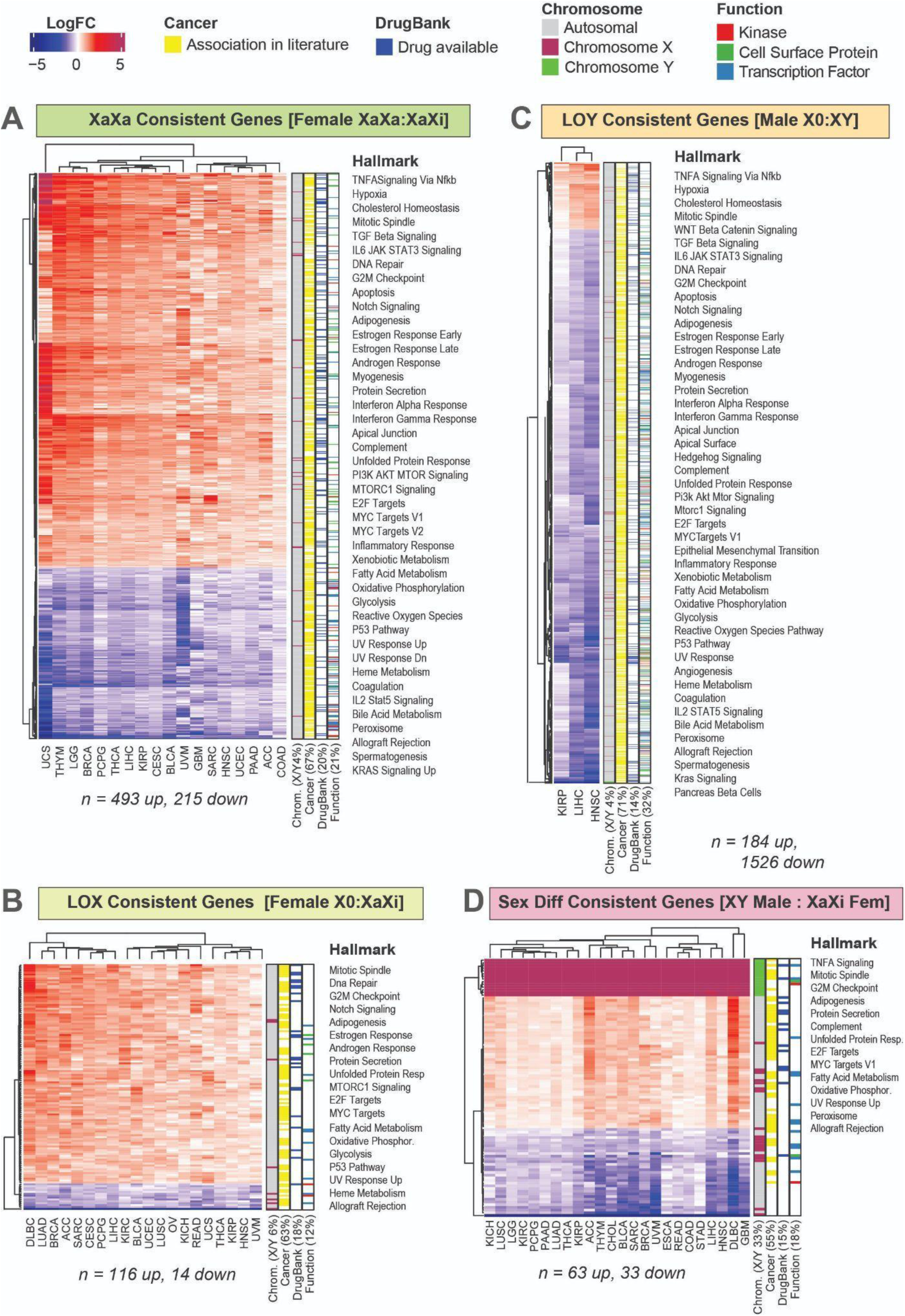
Genes affected by loss of sex chromosomes consistently across many tissue types. Consistent gene expression changes that were in the same direction and statistically significant differential expression (p-value < 0.05) in at least 60% of tissues. Genes changing: **(A)** with reactivation of the X chromosome (XaXa : XaXi) in tumors from female patients, **(B)** with loss of the X chromosome (X0 : XaXi) in tumors from female patients, **(C)** with loss of the Y chromosome (X0 : XY) in tumors from male patients, **(D)** between XY tumors from male patients and XaXi tumors from female patients. Genes are annotated with known functions and associations with hallmarks of cancer.

In tumor samples from female patients, 18 tissues had enough samples to calculate differential expression for maintenance of two X chromosomes (XaXa). Genes changing in XaXa female samples compared to XaXi female samples included 493 genes that consistently increase expression across tumor types and 215 genes that consistently decrease in expression. 20 tissues have samples that could be used to assess X0 compared to XaXi female samples to create a LOX profile. LOX consistent genes are fewer in number but mostly have higher expression in the X0 samples (116 increase in expression and 14 decrease). In tumor samples from male patients, loss of Y could only be assessed in three tissues; within those, 1526 genes have decreased expression in LOY (male X0) samples and nearly 10 times less (184 genes) have increased expression in LOY. In all three profiles, genes consistently changing between groups with different inferred sex chromosome complements include mostly autosomal genes, with only 4-6% of genes being on sex chromosomes. Each profile contained many genes that have been associated with cancer in previously published work (55-71%), have drugs available for possible therapeutic intervention (15-20%), and have functions known to be important in cancer such as kinases, transcription factors, and cell surface proteins (12-32%).

There is a set of genes that are consistently differentially expressed with the loss of a sex chromosome. There are nine genes that are consistently more lowly expressed in both LOX and LOY profiles (*PCM1*, *SPG11*, *HIPK3*, *SENP6*, *CAPN7*, *DYNC1LI2*, *GOLGA4*, *SETD2*, *AKAP17A*). *SETD2* is a histone methyltransferase that is involved in transcription elongation, DNA repair, and gene splicing which is considered to be a tumor suppressor as it is mutated in various cancers including leukemia and renal cancer (Fahey and Davis 2017). *HIPK3* is a serine/threonine kinase involved in cell survival and proliferation; low expression of *HIPK3* has also been observed in renal cancer where it is also thought to act as a tumor suppressor (Xiao et al. 2021). 44 genes are consistently more highly expressed in both LOX and LOY. Many of these genes are part of mitochondrial oxidative phosphorylation complexes (*COX5B*, *COX6B1*, *COX14*, *NDUFB7*, *NDUFB11*, *NDUFS5*, *ATP5F1D*, *ATP5F1E*, *UQCR10*, *SDHAF1*), so this may indicate that tumors that have lost sex chromosomes may undergo metabolic reprogramming. Upregulation of oxidative phosphorylation genes can support resistance to chemotherapies and metastasis (Uslu, Kapan, and Lyakhovich 2024). Several genes upregulated in LOX and LOY are ribosomal proteins (*RPL8*, *RPL18*, *RPL18A*, *RPLP2*, *RPS9*, *MRPL27*, *MRPL52*, *MRPL14*, *MRPS12*) and overexpression of ribosomal genes supports tumorigenesis as increased protein production is needed for cancer progression (Hwang and Denicourt 2024). *SETD2*, related family member *HIPK1*, and mitochondrial ribosome complex genes are also in the XaXa consistent gene profile, indicating that these genes might be related to low *XIST* expression which is common to LOX, LOY, and XaXa states. The LOY profile includes decreased expression of several tumor suppressor genes including *PTEN*, *DLC1*, *RASSF8*, and *TSC2*.

In addition to differentially expressed genes between different SCCs within the same patient sex, we observe a set of genes consistently differentially expressed across tissue types between female and male patients with typical sex chromosome complements. Previous research has identified genes differentially expressed between tumors from female versus male patients (Yuan et al. 2016; Han et al. 2022; Jang et al. 2024), but each patient sex group likely contained tumors that had lost a sex chromosome or reactivated an X chromosome. We identified differentially expressed consistently between patient sexes specifically with typical sex chromosome complements (XX vs XY) (Figure 3). This profile included sex chromosome genes by definition, but also included many autosomal genes for a total of 63 genes that had higher expression in XY male tumors and 33 genes that had higher expression in XaXi female tumors. Genes consistently differentially expressed between sex chromosome complements are associated with hallmarks of cancer, demonstrating the wide array of cancer-related functions that can be in different states based on their X and Y chromosome status.

Mapping the consistent gene profiles by chromosomal location demonstrates that genes differentially expressed in samples with atypical sex chromosome complements are found across the genome (Figure 4). The LOY consistent gene profile includes genes on every chromosome and the XaXa consistent gene profile includes changes in nearly every chromosome (all but chromosome Y and chromosome 18). Chromosome 19 contains a set of genes that have increased expression with *XIST* loss, in both XaXa and LOX. Previous studies suggest that chromosomes 1 and 19 may contain genes that are responsible for maintaining a single active copy of the X chromosome in early development (Migeon, Beer, and Bjornsson 2017). In addition to the clustering of consistently increased genes in chromosome 19 in the XaXa and LOX profiles, genes that are increased in XaXa show a visible cluster in chromosome 1. Smaller clusters are visible in chromosomes 5, 6, 9, 11, and 12, potentially identifying genes necessary for the regulation of X ploidy and *XIST* expression in cancer cells. A portion of the consistent gene profiles include genes that have been shown to interact with *XIST* or have been shown to be targets of the *KDM5D*, one of the chromosome Y marker genes and a transcription factor (see Methods and Table 1), suggest direct regulatory consequences of loss of XIST or Y chromosome expression. Other chromosome Y transcription factors that are expressed in normal adult tissues such as *ZFY* may also directly interact with consistently changing genes, but a full list of targets was only available for *KDM5D* and/or *SRY* in commonly used transcription factor target databases and *SRY* did not have high expression in adult tissues (Figure S1). These genes may be directly interacting with the genes that are affected by having a different sex chromosome complement, while the rest of the consistent gene profiles may be indirect effects.

**Fig. 4.**
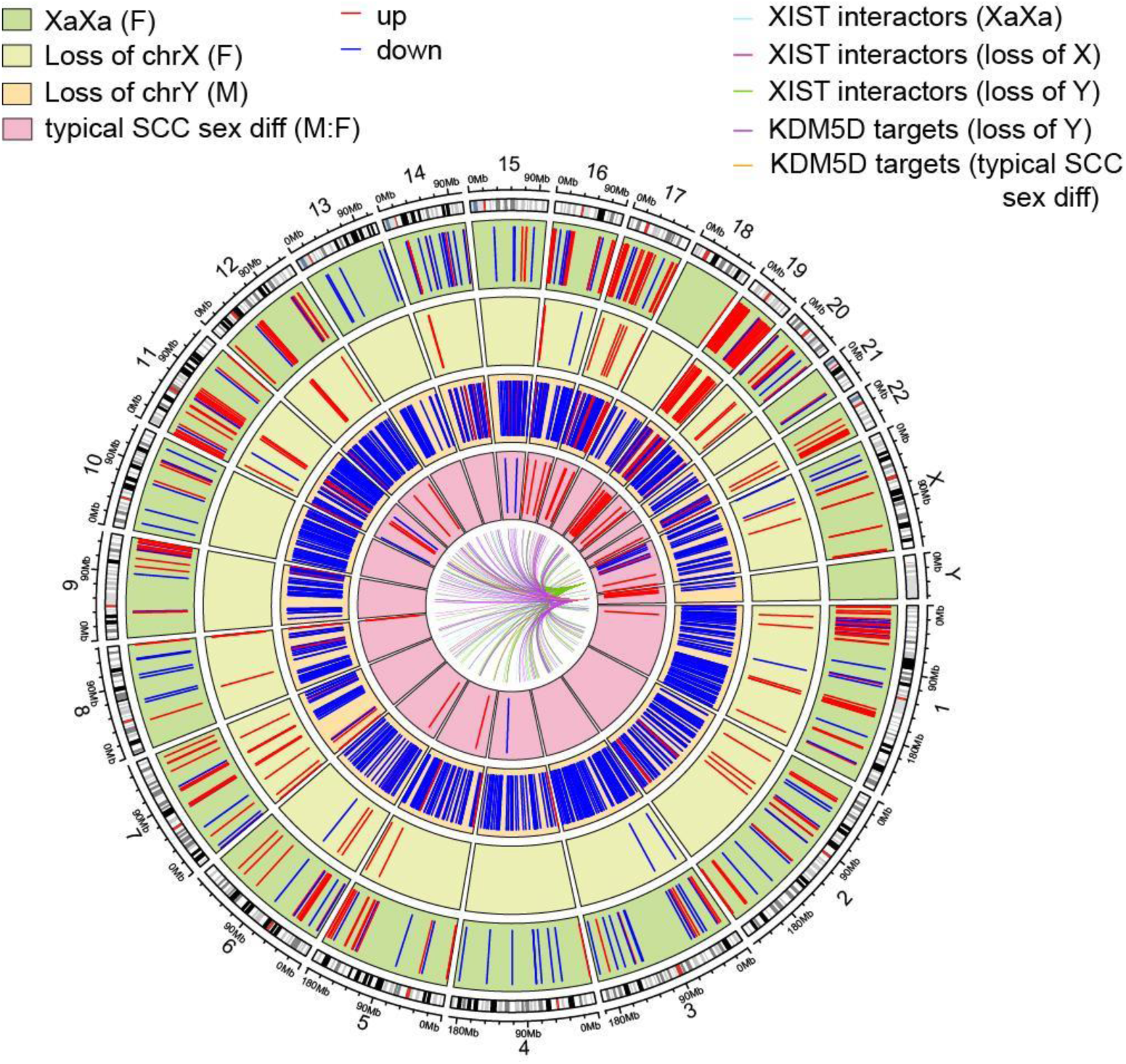
Genomic location of consistent XaXa, LOX, and LOY genes. Circos plot showing the position of genes that are changing significantly over multiple tissues in the genome. Going from outside in, tracks include genes changing on reactivation of the X chromosome in tumors from female patients, genes changing on loss of an X chromosome in female patients, genes changing on loss of the Y chromosome in male patients, and genes changing between XY tumors from male patients and XaXi tumors from female patients. Lines show which consistent genes interact with *XIST* and *KDM5D*.

**Table 1.** Inferred sex chromosome complement of all samples in GTEx, TCGA primary tumors, and CCLE.

**Table 2.** Consistent differentially expressed genes across tissue types in TCGA.

Although we had smaller samples sizes, we further investigated gene expression differences in adult normal tissues (GTEx) and cancer cell lines (CCLE) with evidence of loss or reduced X and Y expression. As sex chromosome ploidy was not available for GTEx, differential gene expression between tissue samples from female individuals was calculated between samples that had expression of *XIST* (> 10 TPM) indicating an XaXi sex chromosome complement and those that did not due to either loss of X or X reactivation (XaXa) (Figure S9, Table 3). Across the 20 tissues that had at least 3 samples with LOX or XaXa, 337 genes were consistently higher in X0/XaXa female tissue samples and 461 genes were consistently lower compared to XaXi female tissue samples (p-value < 0.05 in at least 60% of the tissues and in the same direction). While these genes were identified from normal adult tissues, we observe that many of these genes were involved in cancer-related functions just as the consistently differentially expressed genes in primary tumors were. 36 genes overlapped with the primary tumor female LOX consistent gene profile and 151 genes overlapped with the primary tumor female XaXa consistent gene profile, which is significantly more than would be expected by chance (hypergeometric p-value = 7.22×10^−19^ and 7.48 × 10^−61^, respectively). Overlap of gene expression differences in loss of *XIST* in GTEx with LOX and XaXa in tumors include the ribosomal protein genes (*RPL27A*, *RPL13A*, *RPL18A*, *RPL18*, and *RPS19*), including those involved in mitochondrial function (*MRPL52*, *MRPL55*, and *MRPL28*). These overlapping genes also included *AURKAIP1*, a kinase interaction protein associated with mitochondrial translation, and several genes involved in metabolic processes, including *NDUFB7*, *ATP5F1D*, and *PGLS*. Genes overlapping specifically in XaXa profiles in both GTEx and TCGA included several genes that are part of signaling pathways known to be involved in cancer including *HRAS* in the Ras signaling pathway, *MEF2A* in the p38 pathway, and *SIRT6* and *EP300* in the HDAC pathway. Observing genes changing in groups with different sex chromosome complement that are consistent across tissues with and without cancer increases our confidence in these genes being associated with the sex chromosomes.

**Table 3.** Consistent differentially expressed genes across tissue types in GTEx.

**Table 4.** Samples with unexpected sex chromosome complements in GTEx, TCGA, and CCLE.

Just as there were only three tissues from male patients with which a consistent LOY profile could be made in TCGA tumor tissues, there was only a small number of tissues in GTEx adult normal tissues from male individuals that had enough samples with LOY to measure differential expression– liver and heart. Using the samples with low chromosome Y gene expression, there were 3765 genes that were in the same direction in both tissues and were significant in both (p-value < 0.05 in at least 60% of the tissues which in this case would be both tissues), 1842 increasing with LOY and 1923 decreasing with LOY. Of those, 338 were overlapping with the TCGA tumor LOY consistent gene profile (Figure S9, Table 3). Just like with LOX/XaXa in females in GTEx, a large portion of LOY genes in adult normal tissues were autosomal and associated with cancer-related functions. LOY in both normal adult tissues and primary tumors includes decreased expression of *AKT1*, which can be pro-cancerous when overexpressed in early cancer lesions, but can be tumor suppressive in advanced cancer as *AKT1* is necessary for preserving the integrity of vasculature and decreased expression of *AKT1* can promote metastasis (Alwhaibi et al. 2019; Rao et al. 2017). LOY genes consistently altered in both normal and tumor tissues from males are enriched for membrane trafficking, which is often dysregulated in cancers leading to altered levels of receptor tyrosine kinases, immunosuppressive molecules, and adhesion proteins, all of which are also included in genes with differential expression in LOY male tissues (Figure 3, Figure S9, Table 3).

In cancer cell lines from CCLE, cell lines from different tissues showed dramatically different profiles of differentially expressed genes with *XIST* loss and chromosome Y loss (Figure S10). Applying the same criteria for consistent differential gene expression, the few genes identified were sex chromosome genes and nearly all were marker genes used to identify the sex chromosome complement. Differentially expressed genes between cell lines with and without sex chromosome alterations were identified, but they were present in at most two tissues out of the 12 tissues types that had at least 3 cell lines from female patient tumors in each group (XaXi, X0/XaXa) and 10 tissue types from male patient tumors (XY, X0). Cancer cell lines are highly variable in how they are established and maintained in cell culture. Additionally, the sample size in the CCLE cell lines made it difficult to draw broad conclusions about the effects of loss of typical sex chromosome expression.

### Loss of sex chromosomes can lead to a similar X0 tumor state regardless of the patient sex

While gene expression profiles from human tissues typically separate by reported sex of the individual, loss of sex chromosomes reduces this divide and may bring tumors to a common X0 state. To investigate the outcome of losing sex chromosomes in tumors from female and male patients, gene expression profiles of genes distinguish samples by patient sex and sex chromosome complement were mapped to each other. Analysis of variance was used to determine genes that show differences in expression between at least 2 of 4 sample groups based on patient reported sex and loss of sex chromosomes: tumors from female patients with the expected sex chromosome complement (XaXi), tumors from female patients that had loss of chromosome X (X0), tumors from male patients that had the expected sex chromosome complement (XY), and tumors from male patients that had functional loss of chromosome Y (X0). The top 3000 distinguishing genes (equating to ANOVA p-value < 3.52 × 10^−57^) mapped using partial least squares discriminant analysis show that tumors from female patients that have loss of chromosome X and tumors from male patients that have lost chromosome Y cluster together, compared to tumors that have the expected sex chromosome complement based on the reported sex of the patient (XaXi in female patient samples and XY in male patient samples) (Figure 5, Figure S11). This trend is maintained even when genes from the sex chromosomes are removed from the analysis, demonstrating that there are autosomal genes that distinguish XaXi female tumors from XY male tumors and that that distinction is reduced with LOX in females and LOY in males. This trend is visible through a range of ANOVA significance thresholds; at the most stringent threshold, male samples containing a chromosome Y are the most distinguishable in their gene expression profile even when sex chromosome genes are filtered out (Figure S11). This trend is visible but to a lesser degree in cancer cell lines (Figure S11). Taken together, loss of sex chromosomes causes a shift in both autosomal and sex chromosomal gene expression towards a common X0 state in which tumors from female and male patients are indistinguishable based on gene expression.

**Fig. 5.**
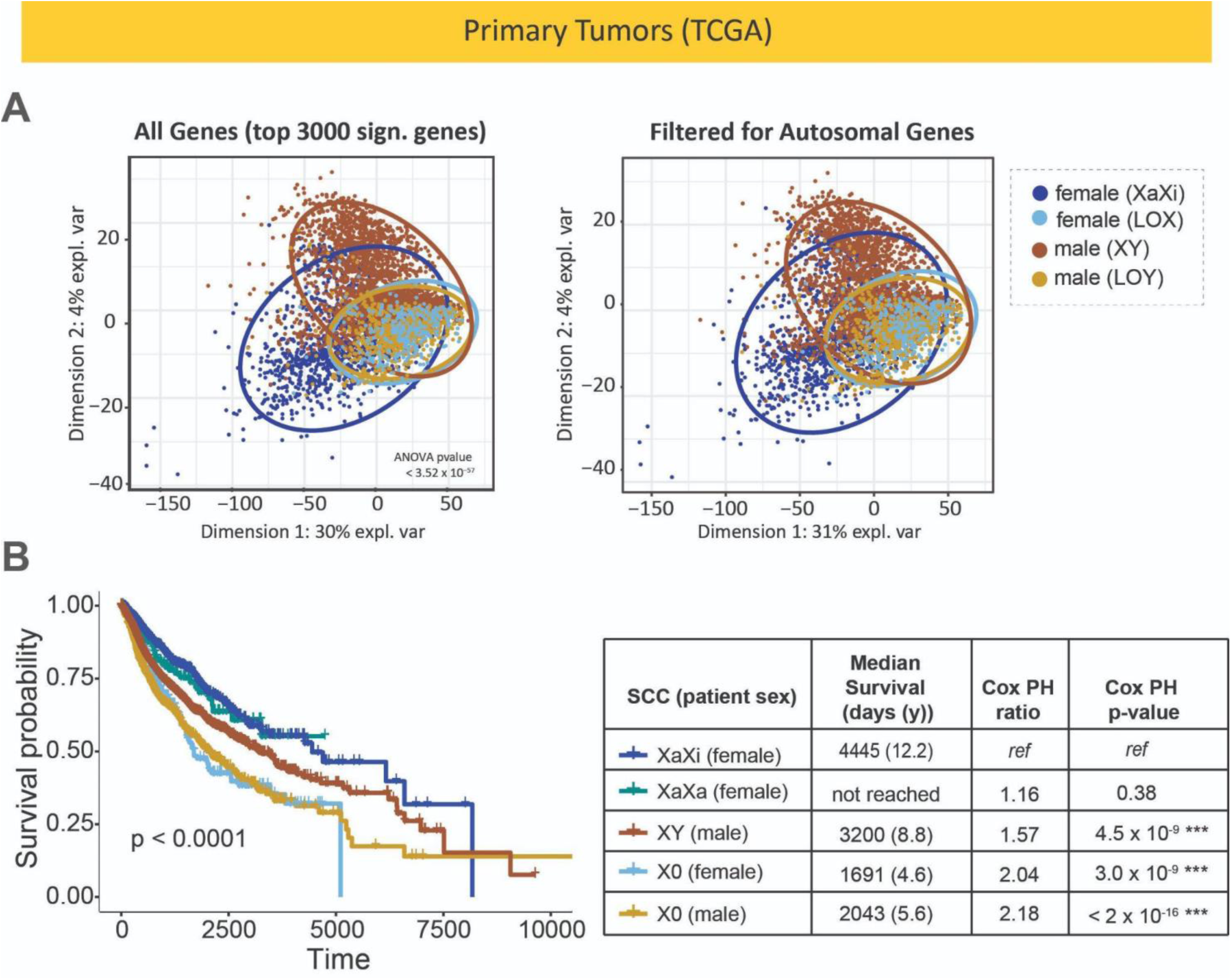
Loss of sex chromosomes can lead to a X0 tumor state with reduced patient survival. **(A)** Top genes that distinguish between tumor samples from TCGA grouped by reported sex and sex chromosome complement (top 3000 by ANOVA p-value between XaXi and X0 samples from female patients and XY and X0 samples from male patients) were mapped using partial least squares discriminant analysis, shown with and without filtering for only the autosomal genes. **(B)** Kaplan-Meyer curves showing the survival of patients with tumors with different sex chromosome complements. Global significance is given on the survival plot; pairwise comparisons between each SCC group are given below.

### Patients with tumors that have lost sex chromosomes show a lower survival

Using clinical information from TCGA (Thorsson et al. 2018), survival probability curves for patients with tumors with various sex chromosomes were compared. Overall, grouping patients by the sex chromosome complement of the tumors isolated from them shows a significant difference in survival (p < 0.0001) (Figure 5D). Male patients with XY tumors had lower survival than female patients with XaXi tumors, but female patients that had tumors that had loss of X and male patients that had tumors that had loss of Y (both X0) had significantly lower survival than both typical XaXi female and XY male groups. Within the X0 groups, survival was equally poor in the female and male patients. Female patients with tumors that had the expected sex chromosome complement (XaXi) had a median survival time of 4445 days (12.2 years), while females with tumors that had lost an X chromosome had a median survival of 1691 days (4.6 years) (Table S1). Male patients with tumors with the expected sex chromosome complement (XY) had a median survival of 3200 days (8.8 years), but male patients with tumors that had lost a Y chromosome had a median survival of only 2043 days (5.6 years). Interestingly, female patients that had XaXa tumors did not have a lower survival probability compared to female patients with XaXi tumors. X0 tumors had higher aneuploidy levels in general which may be due to reduced expression of the *APC* gene which plays a key role in genome stability (Figure S12) (Taylor et al. 2018).

### Differential gene expression profile between patient sexes affected by sex chromosome complement

In cases where different gene expression studies of cancer wish to consider sex as a biological variable, patient sex is typically used to capture that information instead of the sex chromosome complement. This is because patient sex is included in sample metadata but the sex chromosome complement is rarely assessed. However, when the sex chromosome complement is not taken into account, differentially expressed genes between tumors from female versus male patients can be very different. To directly compare the effect of loss of X, loss of Y, and loss of X chromosome inactivation, we sought to examine this against a common background of the same tissue type. Of the three tissue types for which there were enough samples to create a LOX, LOY, and XaXa differential gene expression profile, hepatocellular carcinoma (LIHC) was the only tissue that had a substantial number of differentially expressed genes in all three profiles after accounting for age multiple hypothesis testing (Figure S13). To identify genes that might be missed if the sex chromosome complement was not considered, we looked at genes that had significant differential expression in tumors from female patients with the expected XaXi genotype compared to tumors from male patients that were XY and but that difference was no longer present after loss of X or loss of Y (Figure 6). Genes involved in key signaling pathways in liver cancer often follow the trend of being differentially expressed between XaXi female and XY male tumors but have similar expression in X0 tumors from either patient sex. Genes that had higher expression in XaXi tumors from female LIHC patients were enriched for angiogenesis genes, including *CDH13* (cadherin 13), *THBS2* (thrombospondin-2), and *HSPG2* (heparan sulfate proteoglycan 2). These genes have significantly lower expression in XY male patient samples, and expression as low as XY male samples in X0 samples from both female patients (loss of X) and male patients (loss of Y). Cadherins, thrombospondin, and *HSPG2* have all been shown to have both pro-angiogenic activity and act as inhibitors of angiogenesis in different tumor types (Wen et al. 2025; Carpino et al. 2021; Corbella et al. 2024; Douglass, Goyal, and Iozzo 2015); it is possible that the sex chromosome complement of the tumor may contribute to the complex context that determines how these angiogenesis genes affect tumor cell biology. *TGFBRAP1* and *SMAD5* are known regulators of the TGFβ signaling pathway, which also has a seemingly paradoxical role in liver cancer having shown both pro-and anti-tumorigenic functions (Zhang et al. 2020; Ge et al. 2025; Gungor, Uysal, and Senturk 2022; Tu et al. 2019). *TGFBRAP1* has identified as regulator of TGFβ signaling that governs sensitivity to tyrosine kinase and has increased expression in drug-resistant liver cancer stem cells (Liu et al. 2024), while *TGFA* is ligand that mainly binds to receptor tyrosine kinase EGFR (epidermal growth factor receptor) and promotes tumor cell proliferation through the EGFR/RAS/MAPK signaling pathway. Sex differences in TGFβ signaling and EGFR signaling have been demonstrated in other cancers and other conditions (James 2016; Ziller et al. 2020; Frydecka et al. 2015; Huang et al. 2022), suggesting sex chromosome differences might explain some of the paradoxical nature of interacting signaling pathways leading to sex differences in cancer biology.

**Fig. 6.**
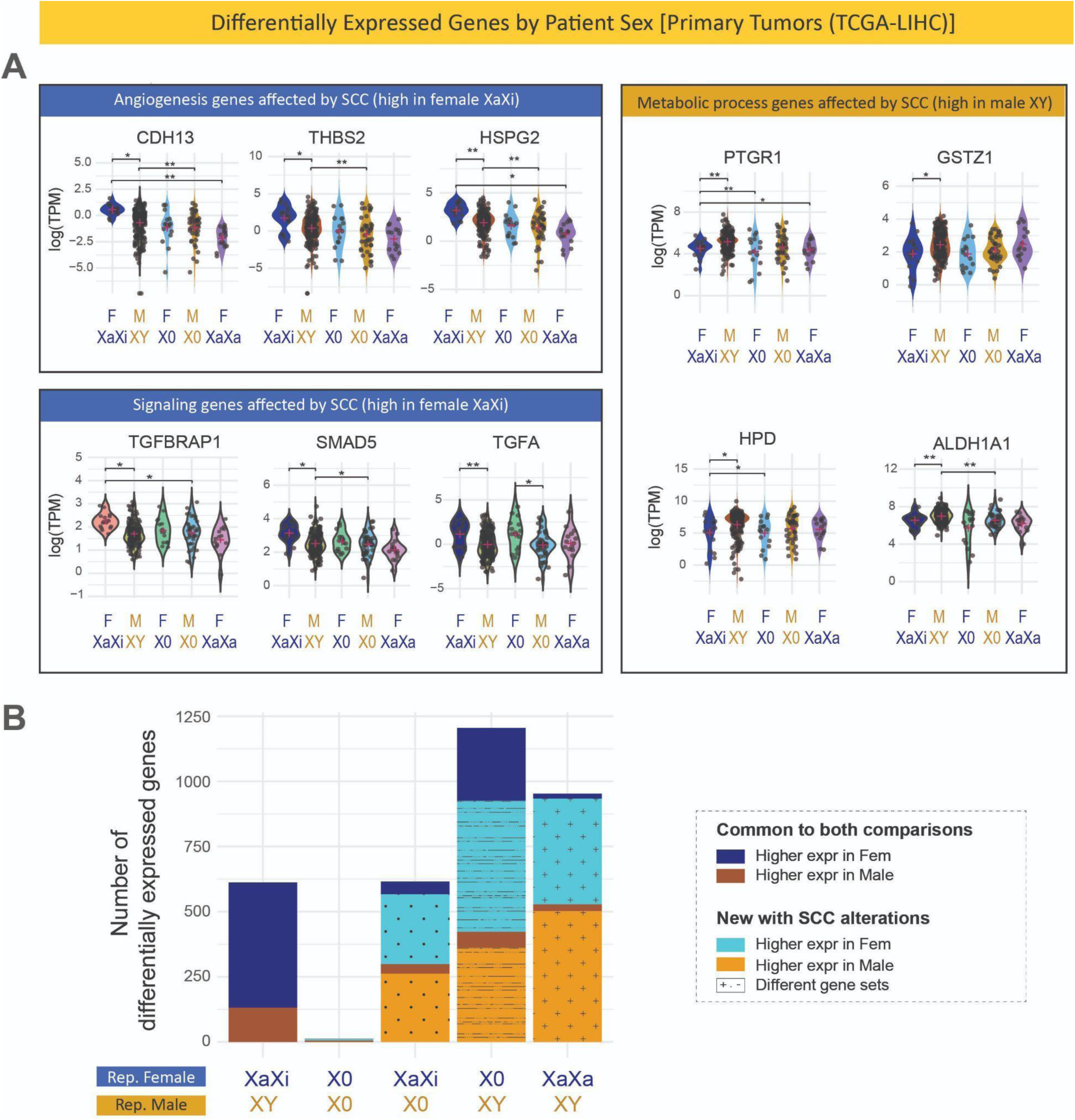
Loss of sex chromosomes reduces patient sex differences in gene expression in primary tumors. (**A**) Genes involved in enriched gene functions relevant to cancer that show sex differences between tumors with typical sex chromosome complements that are no longer showing sex differences when sex chromosome complements are not typical. Comparisons marked with a single asterisk (*) indicate differential expression adjusted p-values less than 0.05 and those marked with double asterisks (**) indicate adjusted p-values less than 0.01. (**B**) The number of significantly differentially expressed genes between liver tumor samples (TCGA-LIHC) from female versus male patients. The number of differentially expressed genes between samples with the expected sex chromosome complements is shown as reference (adjusted p-value < 0.05, abs(log_2_ foldchange) > 1) and the analysis is repeated with loss of chromosome X in female patient tumors, loss of chromosome Y in male patient tumors, both, or activation of the second X chromosome in female patient tumors. Genes are separated by colors to indicate which patient sex had higher expression, whether the genes were autosomal or in chromosomes X or Y, and whether they are shared with the reference comparison with typical sex chromosome complements or are unique to comparisons where sex chromosome complements are not typical.

Genes that had significantly higher expression in XY tumors from male patients than in XaXi tumors from female patients were enriched for genes involved in metabolic processes. *PTGR1* (prostaglandin reductase 1), *GSTZ1* (glutathione S-transferase zeta 1), and *HPD* (hydroxyphenylpyruvate dioxygenase), and *ALDH1A1* (aldehyde dehydrogenase 1) are all enzymes involved in metabolic functions that have been shown to play a role in cancer and have been observed to also play both pro- and anti-tumorigenic effects (Yue et al. 2022). Both *PTGR1* and *GSTZ1* are associated with the NRF2 (erythroid 2-related factor 2) pathway which is a master regulator of detoxification and antioxidant response which lead to metabolic changes that are a hallmark of cancer (J. Li et al. 2019). *PTGR1* is involved in the catabolism of eicosanoids and lipid peroxidation; it has been shown to be involved in the response to oxidative stress being a high proliferation rate and evasion of cell death in liver cancer tumors (Sánchez-Rodríguez et al. 2017; X. Wang et al. 2021). *GSTZ1* is involved in phenylalanine/tyrosine catabolism. *GSTZ1* deficiency causes overactivation of the NRF2 antioxidant response pathway and promotes progression in liver cancer (J. Li et al. 2019). *ALDH1A1* is associated with cancer cell stemness in many cancers. *ALDH1A1* levels have been associated with survival in liver cancer (Tanaka et al. 2015; Gehlot et al. 2016; Pommergaard et al. 2022). In all of these examples, gene expression is significantly higher in XY tumors from male patients, but is lower in tumors from female patients with and without loss of sex chromosomes. As sex differences in metabolism have been reported during development, into adulthood, and in both health and disease, the overexpression of metabolism genes in the presence of a Y chromosome demonstrates possible contributors to mechanisms underlying sex differences in metabolic reprogramming that occurs in tumors.

To determine how common this trend was in liver cancer, we examined how many differentially expressed genes were found between samples from female patients versus samples from male patients and how those differentially expressed gene lists were altered with loss or reactivation of sex chromosomes (Figure 6). Within liver cancer patient samples, 612 genes showed differential gene expression by patient sex when the samples had the expected sex chromosome complement (XaXi female patient samples versus XY samples from male patients), 481 with higher expression in XaXi female tumors and 117 in XY male tumors (multiple hypothesis testing corrected p-value < 0.05, fold change > 2). The majority of those sex differentially expressed genes (584 of 612) were autosomal. Genes with significantly higher expression in female patient samples were enriched for cellular adhesion, angiogenesis, and cell morphogenesis, while genes significantly higher in male patient samples were enriched for metabolic processes (Table 5). The vast majority (all but 12 of these genes) are no longer differentially expressed between female and male patient tumors in cases where the females had lost chromosome X and males had lost chromosome Y. Comparing tumors from female patients with tumors from male patients when only one group had atypical sex chromosome complements shows differential expression of a host of new genes (Figure 6). Genes that were differentially expressed with sex chromosome complement alterations were associated with many functions and pathways associated with cancer, including cell cycle, oxidative phosphorylation, intracellular transport, and translation (Table 5). Failing to take sex chromosome complement into account when looking for patient sex differences in cancer can lead to vastly different results from differential expression studies.

**Table 5.** Differential expression results between patient sex in liver cancer (TCGA-LIHC) with different sex chromosome alterations.

## Discussion

The results of this study show that the reported sex of the patient does not always predict the functional sex chromosome complement of tumors and cancer cell lines. Sex chromosome marker gene analysis identified loss of sex chromosomes and loss X chromosome in activation in a large portion of tissues across tissue types. Sex chromosome gene expression analysis also identified cases where samples from male patients appear to have two X chromosomes as noted by *XIST* expression, particularly in testicular germ cell tumors. *XIST* expression and increase in X chromosomes in tumors from male patients has been described as a biomarker for this tumor type, having been associated to demethylation of the *XIST* promoter and methylation of the androgen receptor (Weakley et al. 2011; Kawakami et al. 2004; Lobo et al. 2019; Kawakami et al. 2003; Looijenga et al. 1997). Cases where reported sex of the patient was likely inaccurately annotated in patient records or laboratory logs were also identified; samples that had expression of chromosome Y genes and no *XIST* expression are highly unlikely to have come from a female patient. In cancer cell lines, a proportion of cell lines were shown to have lost sex chromosome gene expression in nearly every tissue type. In some cases where the reported sex of the patient from which the cancer cell line was derived was not annotated, the sex chromosome complement can be used in place of patient sex. Taken together, it is critical to check expression of sex chromosome genes in genomics datasets to assess the sex chromosome complement early in analysis workflows as this has a large effect on overall gene expression in the tissue.

Reduction in *XIST* and Y-linked gene expression in tumors and cell lines could be due to different rates of sex chromosome loss or maintenance of two X chromosomes in different cell types in the primary tumors, as tumors can be quite heterogeneous in the cell types present in the tumor sample and the progression of cancer can be different in various tumor cell subpopulations. Previous studies demonstrate that primary tumors from TCGA had high variability in the cell types contributing to bulk gene expression signatures (Tiong, Luzhbin, and Yeang 2024; Revkov et al. 2023; Xue et al. 2024; Newman et al. 2019), so it is possible that specific cells have lost sex chromosomes or reactivated sex chromosomes while others have not and the average expression of these genes is lower in samples where more cells have lost sex chromosomes. A recent study reported a high correlation between expression of chromosome Y genes and chromosome Y copy number at the single cell level in epithelial cancer types and determined that cells that have lost chromosome Y gene expression also show downregulation of a number of pathways related to cancer phenotypes (Chen et al. 2025). The wide range of sex chromosome gene expression in primary tumors may also reflect a range of tumor purity in different tumor samples (Aran, Sirota, and Butte 2015; Revkov et al. 2023). Examining loss of sex chromosomes in single cell data sets will allow for identification of sex chromosome changes in specific cell populations in health and disease states.

A potential limitation of our method to infer sex chromosome complement using the expression of specific sex chromosome genes is that expression of these genes is lower overall in specific non-cancerous adult tissues. Most brain tissues and whole blood have a lower average expression of *XIST* and chromosome Y genes than other non-cancerous adult tissues. This may be an artifact of the relative size and shape of cells in specific tissues that may dilute gene expression measurements, such as brain cells having a long axon. The range of sex chromosome gene expression in tumors and cancer cell lines from the brain and blood don’t appear to have this reduction; further study is required to understand how expression thresholds for *XIST* and chromosome Y genes may need to be adjusted for specific tissue and cell types.

This work provides a resource for researchers using three common genomics data sets (GTEx, TCGA, and CCLE) to look up the inferred sex chromosome complement in the tissues they are studying. For primary tumor tissues, loss of sex chromosome gene expression or inferred sex chromosome complements can be used as a covariate when analyzing changes in clinical features in order to make results more robust to sex differences. For cell lines used as preclinical models in cancer research, cancer cell lines from male patients that have lost chromosome Y gene expression may serve as better models for primary tumors that have lost chromosome Y in male patients. For example, loss of chromosome Y in bladder cancer was associated with a lower prognosis and evasion of the immune system (Abdel-Hafiz et al. 2023) and bladder cancer cell lines derived from male patients with and without chromosome Y gene expression can be found in the CCLE (BLCA, Figure 2B, Figure S3), so the cell lines that have lost chromosome Y gene expression could be a better model of tumors that have lost chromosome Y. Similarly, female patient cell lines that have lost *XIST* expression may be a better model for primary tumors that have lost an X chromosome or have impaired X chromosome inactivation. Loss of *XIST* expression in ovarian cancers has been linked to increased cancer cell stemness and poorer patient survival (Naciri et al. 2024), so perhaps comparing ovarian cancer cell lines that have *XIST* expression to those that have lost *XIST* expression (OV, Figure 2B) could be useful in figuring out the genes contributing to increased stemness. Likewise, as cancer stem cells might undergo loss of X chromosome inactivation as induced pluripotent stem cells and embryonic stem cells from female donors do (Brenes et al. 2021; Cloutier et al. 2022), XaXa cell lines might be a better model system for studies of this cancer cell type. Additionally, repeating experiments in cell lines that have different sex chromosome complements might allow researchers to more faithfully recapitulate sex differences in tumor cell biology that can contribute to the understanding of sex differences in clinical features of cancer.

The technique of using specific sex chromosome genes to infer sex chromosome complement can be easily implemented for any transcriptomic data sample in which the expression of these marker genes is measured. In a previous study focused on the loss of chromosome Y in primary tumors from TCGA, Qi et al. used DNA copy number assessed using whole exome sequencing (WES) in tumors compared to matched blood a control as a primary method to identify loss of Y, and presented chromosome Y gene expression-based methods in which chromosome Y gene expression was compared to housekeeping genes as an alternative for when WES or normal controls were not available (Qi et al. 2023). The authors stated that gene expression-based assessment of loss of Y gave nearly identical results as loss of chromosome Y DNA copy number and that their DNA copy number-based rate of loss of Y is likely conservative and underestimated as most controls in TCGA are peripheral blood samples which may themselves be subject to age-related somatic loss of Y. This would make Y copy number in tumor samples appear to be higher. As loss of Y calls are based on differences in fraction of somatic loss of Y in the control versus tumor, no loss of Y is identified when those fractions are roughly equal. We do confirm in our GTEx analysis that whole blood does have a different range of sex chromosome gene expression, so using this as a control may have unintended effects. Our method opts for a simpler thresholding approach in which no comparisons are made to different tissue types which can have different levels of expression of sex chromosome genes, to housekeeping genes which may change in response to cancer or treatment (Irie et al. 2023; J. Wang et al. 2023; Khimani et al. 2005), or between DNA and RNA based measurements taken from adjacent tumor tissue samples which may be quite different in highly heterogeneous tumor types. Sex chromosome gene expression is commonly captured in modern sequencing-based platforms and is downstream of any transcriptional repression that might be in place. While protein level measurements would be further downstream of any translational repression, protein level measurements are impossible for *XIST* since it is not translated and proteomics measurements are less common and less robust for capturing chromosome Y gene expression.

Simple approaches to study sex chromosomes in genomics data will encourage the addition of sex chromosome complement inference in genomics workflows and thus help to identify and characterize sex differences in biology in health and disease. The recommendation for analyzing the sex chromosome complement in tissues would be to obtain the reported sex of the patient and then look at the expression of *XIST* and the chromosome Y marker genes. For tissue samples from female patients, if *XIST* expression is not detected, DNA sequencing can be used to determine X chromosome ploidy to distinguish between loss of X or loss of X chromosome inactivation. For tissue samples from male patients, check for expression of at least two chromosome Y marker genes. It is important that more studies include identification of the sex chromosome complement, not just patient sex, as sex chromosomes are an important molecular determinant of sex differences in cancer and can have a profound impact on the global gene expression landscape of tumor tissues.

## Materials and Methods

### Gene expression data

Gene expression data and sample metadata for normal, non-cancerous tissues were obtained from the Genotype-Tissue Expression (GTEx) consortium (version 8) (GTEx Consortium 2013; Carithers and Moore 2015; GTEx Consortium 2020). Bulk tissue expression data was downloaded as transcripts per million, after mapping, data conversion, and quantification done by the GTEx consortium. Open-access gene expression data for human primary tumors were obtained from The Cancer Genome Atlas (TCGA) (Cancer Genome Atlas Research Network et al. 2013) using the Genomic Data Commons application programming interface (GDC API) through the TCGAbiolinks package (version 2.32.0) in R (version 4.4.0). The data categories indicated were “Transcriptome Profiling” and “Clinical”, experimental strategy was “RNA-Seq”, data type was “Gene Expression Quantification”, workflow type was “STAR - Counts” using the hg38 human reference genome, and quantification method was “tpm_unstranded”. For each of the selected TCGA studies, metadata was extracted using case IDs and retrieved using the GDC API using the GDCquery function with appropriate filters for the project ID. Relevant sample metadata including clinical data, sample types, sex, vital status, survival times, follow-up days, and age at index were extracted for use in specific analysis. Gene expression data and sample metadata for the cancer cell lines were obtained from the Cancer Cell Line Encyclopedia (CCLE) (2019 release) using the web portal (https://depmap.org/portal) (Ghandi et al. 2019; Barretina et al. 2012; Tsherniak et al. 2017). RNA sequencing data quantified as transcripts per million and accompanying sample metadata were downloaded for analysis. Processing of genomics datasets was computed on the high performance computing cluster Sol at Arizona State University (Jennewein et al. 2023).

### Selection of genes used for inferring sex chromosome complement

We chose a set of X-linked and Y-linked genes to use to infer a functional sex chromosome complement (SCC) based on their biology, expression level, and expression pattern across adult tissues using the GTEx data set as reference (Figure S1A). Characterization of the X chromosome in tissue samples was based on the expression of *XIST* as its expression is triggered by the presence of two (or more) copies of chromosome X (Sahakyan, Yang, and Plath 2018). In an XX individual, loss of *XIST* expression could be due to loss of one of the X chromosomes or due to the transcriptional repression of *XIST* which would allow both X chromosomes to remain active. We utilized sex chromosome DNA ploidy information when available to distinguish between (a) loss of XIST due to loss of chromosome X (LOX) and (b) loss of *XIST* expression with maintenance of two copies of the X chromosome, implying two active copies of the X chromosome (XaXa). Characterization of the Y chromosome in tissue samples was based on the expression of genes on the Y chromosome, specifically protein-coding genes on chromosome Y that are outside of the pseudoautosomal regions that are expressed in adult tissues (not just in early in development and/or specifically in gonadal tissues) (Godfrey et al. 2020). Loss of expression of chromosome Y genes has been used to characterize loss of chromosome Y in tumor samples in previous studies (Chen et al. 2025; Qi et al. 2023). Given the expression thresholding scheme implemented (see below), only chromosome Y protein-coding genes whose distribution of expression in samples from male individuals was mostly above our expression threshold (at least 10 transcripts per million), leaving *DDX3Y*, *EIF1AY*, *KDM5D*, *RPS4Y1*, *USP9Y*, *ZFY*, and *UTY* as marker genes (Figure S1A). This is the same set of chromosome Y genes chosen to infer functional loss of chromosome Y in previous studies (Qi et al. 2023).

*Gene expression thresholding scheme*. For all of the genes chosen to infer the sex chromosome complement, we employed a thresholding scheme: greater than or equal to 10 transcripts per million (TPM) RNA sequencing reads defined a gene as being expressed, between 1 and 10 TPM as having low expression, and less than or equal to 1 TPM being not expressed. For the X chromosome, the functional sex chromosome complement was deemed to be XX if *XIST* expression was greater than or equal to 10 TPM. For the Y chromosome, while high overall correlation was observed between the expression of selected chromosome Y marker genes (Figure 1, Figure S1B), expression of at least two of these genes was required to mark the presence of an active chromosome Y. This reduces the rate of false positives but still remains flexible to allow for variability in Y chromosome gene expression in different organs and between different individuals.

### Validation of sex chromosome marker genes at the protein level

Tissue profile data from quantitative immunofluorescence data from the Human Protein Atlas was downloaded and filtered for chromosome Y marker genes. For each chromosome Y gene, protein abundance was expressed in categories: Not detected, Low, Medium, and High. Of the chromosome Y genes observed to have high expression in normal tissues, *DDX3Y*, *EIF1AY*, *RPS4Y1, UTY* and *ZFY* were shown to have at least low expression in many tissue types using quantitative immunofluorescence as indicated in the Human Protein Atlas (Figure S2). Quantitative mass spectrometry-based proteomics data was obtained in order to directly compare relative protein abundance to gene expression. Quantitative mass spectrometry data for normal, non-cancerous tissues was obtained from a publication from the GTEx consortium (Jiang et al. 2020), (Supplemental Tables S1 (sheets A and B) and S2 (Sheets D and F)). Quantitative proteomics data for TCGA was obtained from the Clinical Proteomic Tumor Analysis Consortium (CPTAC) using the Proteomic Data Commons (PDC) web portal as previously described (Y. Li et al. 2023): primary human tumors (Proteome_BCM_GENCODE_v34_harmonized_v1) and matching RNA sequencing data (ALL.rsem_genes_tpm.txt) . Quantitative proteomics data for cancer cell lines in the CCLE (Nusinow et al. 2020) was obtained from the Gygi lab website (https://gygi.hms.harvard.edu/publications/ccle.html). Only three chromosome Y genes had protein abundance above background levels: DDX3Y, EIF4AY, and RPS4Y1 and all showed correlation between gene expression and relative protein abundance (Figure S5). Since *XIST* is a non-coding transcript, we would not be able to perform this protein level comparison. Correlated protein abundance increases our confidence and ability to use these chromosome Y genes to infer the presence of an active Y chromosome broadly across many tissues.

### Differential gene expression profiles

Differential gene expression was assessed using the limma-voom pipeline (Law et al. 2014; Ritchie et al. 2015), both for differential expression between samples from male and female patients and for differential expression within the same patient sex with different sex chromosome complements. For consistent gene expression signatures across tumor tissue types, genes that were considered consistent were defined as those that are changing in the same direction in all tissues and have a differential expression p-value of less than 0.05 in at least 60% of the tissues in which differential expression could be performed (at least 3 samples in each of the groups being compared). The LOX consistent gene profile included genes differentially differentially expressed between samples from female patients that had inferred sex chromosome complement (SCC) of XaXi (XX DNA ploidy and expression of *XIST*) versus samples from female patients that had an inferred SCC of X0 (X DNA ploidy and loss of *XIST* expression). The XaXa consistent gene profile compared samples from female patients that had an inferred SCC of XaXi versus XaXa (XX DNA ploidy and loss of *XIST* expression). Overlap between LOX, XaXa, and LOY consistent gene profiles was visualized using an upset plot (`upset` function from the `UpSetR` package in R v1.4.0). The typical sex differences gene profile was included genes differentially differentially expressed between samples from female patients that had inferred sex chromosome complement (SCC) of XaXi (XX DNA ploidy and expression of *XIST*) versus samples from male patients that had an inferred SCC of XY (wildtype levels of chromosome Y at the DNA level and loss of *XIST* expression). DNA level measurements were taken from previously published work (Qi et al. 2023).

### Enrichment analysis

Genes consistently differentially expressed with LOX, XaXa, LOY, and by patient sex were annotated using several resources (Figure 3). Whether the gene was in a sex chromosome or autosome was attained from human reference genome annotation downloaded from the UCSC Table Browser (Perez et al. 2025). Annotation/enrichment analysis was completed using Metascape (http://metascape.org) (Zhou et al. 2019). Genes were annotated for whether they were associated with cancer if the word ‘cancer’ or ‘tumor’ occurred in the ‘Disease & Drugs (ChatGPT)’ or ‘Protein Functions (ChatGPT)’ columns in the Metascape annotation results. Genes were marked as having drug targets available if the ‘Drug (DrugBank)’ column in the Metascape results were not blank or ‘None’. Genes involved in pathways involved in the hallmarks of cancer were labeled using enrichment results in the ‘Hallmark Gene Sets’ column in Metascape results. Kinases were marked as those found in KinBase (http://kinase.com/) (Manning et al. 2002). Transcription factors were marked as those found in the University of Toronto transcription factor database (https://humantfs.ccbr.utoronto.ca/) (Lambert et al. 2018). Cell surface genes that could be targeted by drugs and antibodies were marked as those found in the cell surfaceome database (downloaded from supplementary tables in the publication about the resource (da Cunha et al. 2009)).

### Circos plot

A circos plot showing the genomic location of genes consistently differentially expressed with LOX, LOY, and XaXa in multiple tissues was constructed using the `circlize` package in R (v0.4.16). Cytoband and coding sequence coordinates for genes in each consistent gene list were downloaded from UCSC Genome Browser (Perez et al. 2025). Genes in each profile were added to three different tracks using the `circos.genomicRect` function. Genes in the consistent gene profiles that interact with *XIST* were marked as those found in the *XIST* interactome (Minajigi et al. 2015). Predicted targets of *KDM5D* (transcription factor on chromosome Y) that were in the consistent gene profiles were marked according to targets found in the Harmonize 3.0 project (https://maayanlab.cloud/Harmonizome/) (Rouillard et al. 2016; Diamant et al. 2025) and the Institute for Systems Biology’s transcription factor database tool (https://tfbsdb.systemsbiology.net/) (Plaisier et al. 2016). These resources only contained target lists for two transcription factors on chromosome Y, *KDM5D* and *SRY*. As *SRY* is known to be expressed only in early stages of development and shown to have very low expression in normal adult tissues (Figure S1), only transcription targets of *KDM5D* were included in the center of the circos plot.

### Analysis of variance (ANOVA)

Inferred sex chromosome complements were used to do an analysis of variance between 4 groups: samples from female patients with *XIST* expressed, samples from female patients without *XIST* expressed, samples from male patients with presence of chromosome Y, and patients from male patients without presence of chromosome Y. The small number of samples with unexpected sex chromosome complements (samples from female patients with chromosome Y genes expressed and samples from male patients with *XIST* expressed) were removed. An ANOVA p-value between these four groups was calculated using the `aov` function (`stats` package v4_4.2.1, R v4.2.1). To account for the differences in overall p-value significance due to differences in the number of samples being compared, top significant genes were selected and mapped using partial least squares discriminant analysis (`plsda` and `plotIndiv` functions in `mixOmics` package v6.20.0, R v4.2.1).

### Survival analysis

Clinical data for TCGA was obtained from previously published work (Thorsson et al. 2018). Tumors were labelled and separated based on patient sex and sex chromosome complement and Kaplan-Meyer curves with a global statistical significance of separation of survival curves were produced (`Surv`, `survfit`, and `ggsurvplot` functions from the `survminer` package v0.5.0 and the `survival` package v3.8-3). Multiple hypothesis testing corrected p-values for that statistical significance of separation between each pair of survival curves was calculated (`pairwise_survdiff` function). To compare sex chromosome loss to loss of other chromosomes, aneuploidy scores and annotation of chromosome deletions were obtained from previously published work (Taylor et al. 2018).

## Acknowledgments

Exploratory research that formed the basis of this work was conducted as part of a Course-based Research Experience conducted at Arizona State University (BIO 498/598). We acknowledge all students of this course for their contribution to the discussion that led to the analysis and resources featured in this publication. The authors acknowledge Research Computing at Arizona State University for providing HPC and storage resources that have contributed to the research results reported within this paper.

## Author contributions

Conceptualization: MAW, SBP, KHB

Methodology: MAW, SBP, RP, SCM, MF, TA, MV

Investigation: MAW, SBP

Visualization: MAW, SBP, RP, MF, ML, TA, MS, IR, JH, AM, JDR

Supervision: MAW, SBP

Writing—original draft: MAW, SBP, MF, ML Writing—review & editing: MAW, SBP

## Data and materials availability

All data used in this work are publicly available in data repositories or provided with previous publications. Descriptions of how to attain the data used in this work are described in the Materials and Methods. Code used to do the analysis in this work can be found in this repository: https://github.com/SexChrLab/FunctionalXandY

