## Supplementary Figures for "Reproducible autosomal gene expression changes with loss of typical X and Y complement across tumor types"

**Fig. S1. Expression and correlation in gene expression of sex chromosome genes used for sex chromosome complement inference**. **(A)** Violin plots showing the distribution of sex chromosome gene expression in all normal, non-cancerous tissues from GTEx by reported sex of the individual. Expression values are shown as log-transformed transcripts per million (TPM). Genes chosen for sex chromosome complement inference are in the top row (bold) based on their distinct expression by sex and having greater than log-transformed expression greater than 1 on average in these adult tissues. **(B)** Correlation coefficients between expression of chosen sex chromosome genes. All comparisons with a coefficient greater than 0.5 are statistically significant.

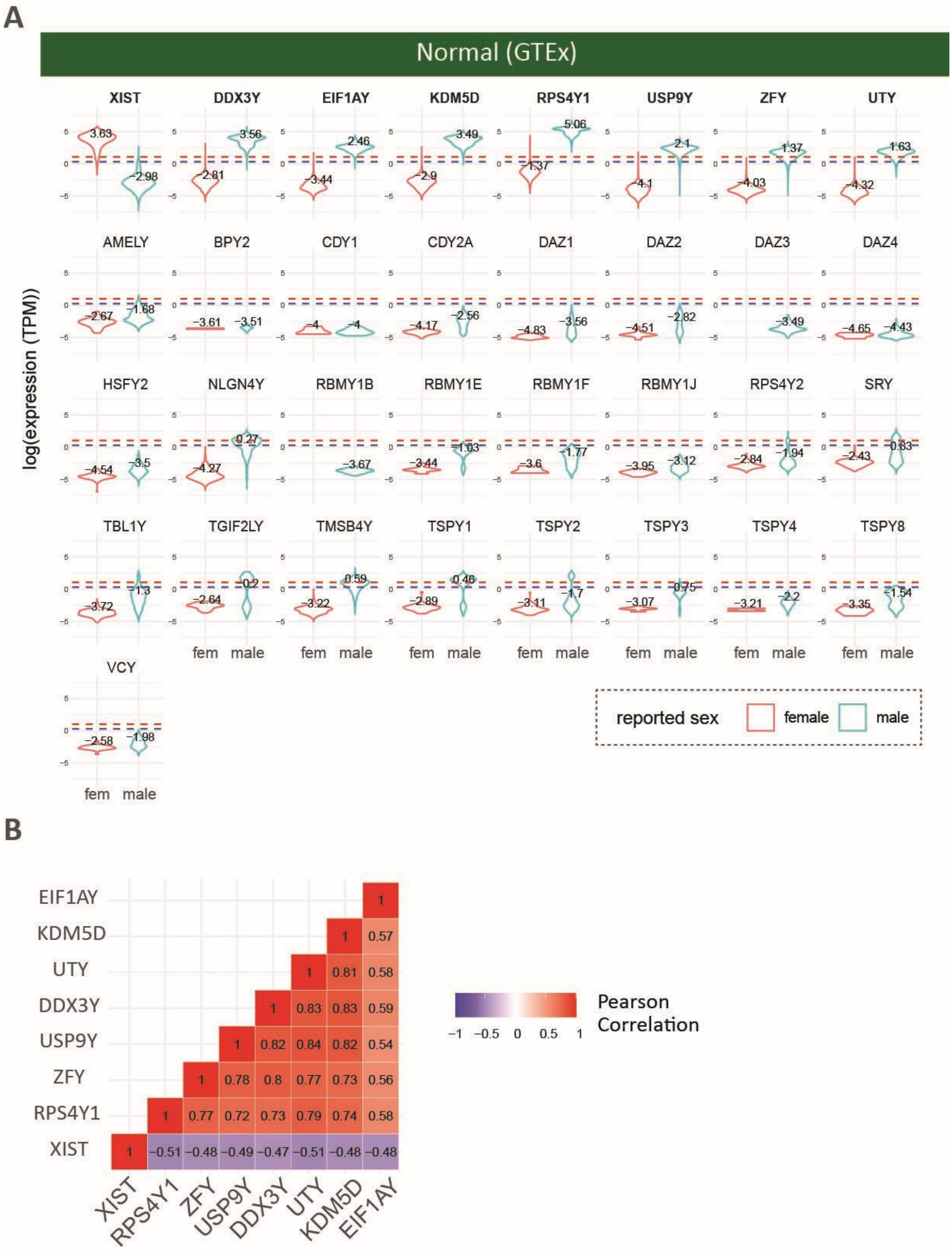

A
Normal (GTEx)
XIST
DDX3Y
EIF1AY
KDM5D
RPS4Y1
USP9Y
ZFY
UTY
AMELY
BPY2
CDY1
CDY2A
DAZ1
DAZ2
DAZ3
DAZ4
HSFY2
NLGN4Y
RBMY1B
RBMY1E
RBMY1F
RBMY1J
RPS4Y2
SRY
TBL1Y
TGIF2LY
TMSB4Y
TSPY1
TSPY2
TSPY3
TSPY4
TSPY8
VCY
log(expression (TPM))
fem
male
reported sex
female
male
B
EIF1AY
KDM5D
UTY
DDX3Y
USP9Y
ZFY
RPS4Y1
XIST
Pearson
Correlation

**Fig. S2. Chromosome Y genes detected in Human Protein Atlas.** Quantitative immunofluorescence based quantification of chromosome Y genes listed in the Human Protein Atlas tissue profile. Each dot represents measurement of each gene (protein product) in a specific tissue type and protein abundance was reported as “High”, “Medium”, “Low”, or “Not Detected”. All genes that have quantification information have at least low expression in multiple tissues.

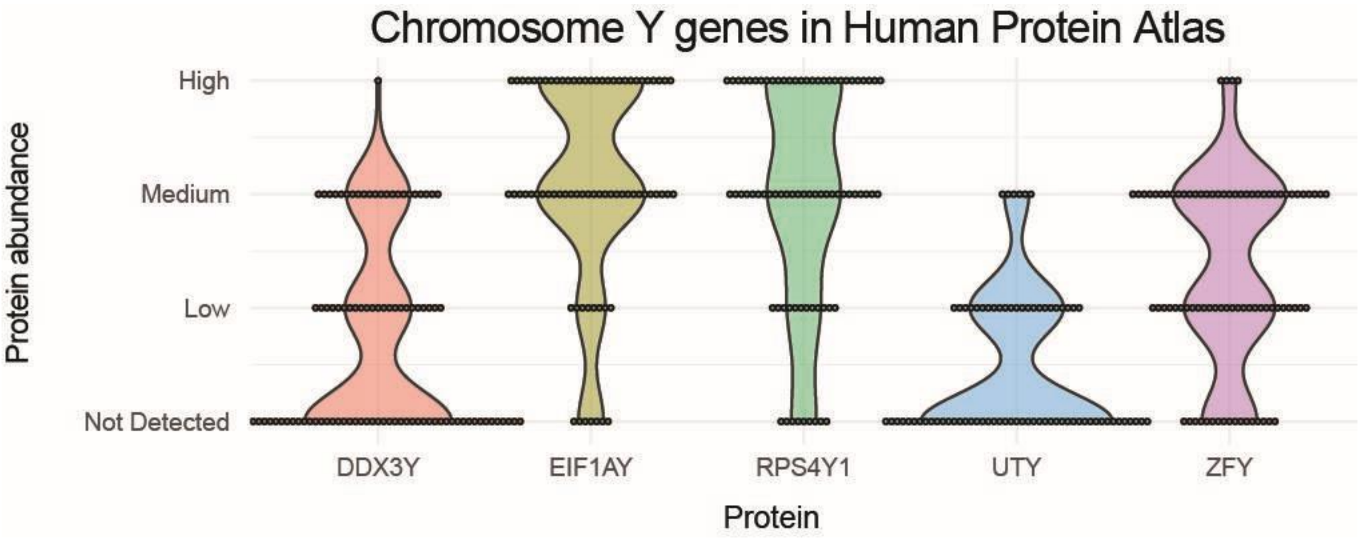

**Fig. S3. Proportion of sex chromosome complements by reported sex of the individual.** For samples reported female and males from GTEx, TCGA, and CCLE, the proportion and number of each inferred sex chromosome complement identified based on sex chromosome marker gene expression is labelled. For some SCCs that were identified at less than 5% of the samples, only the count is provided.

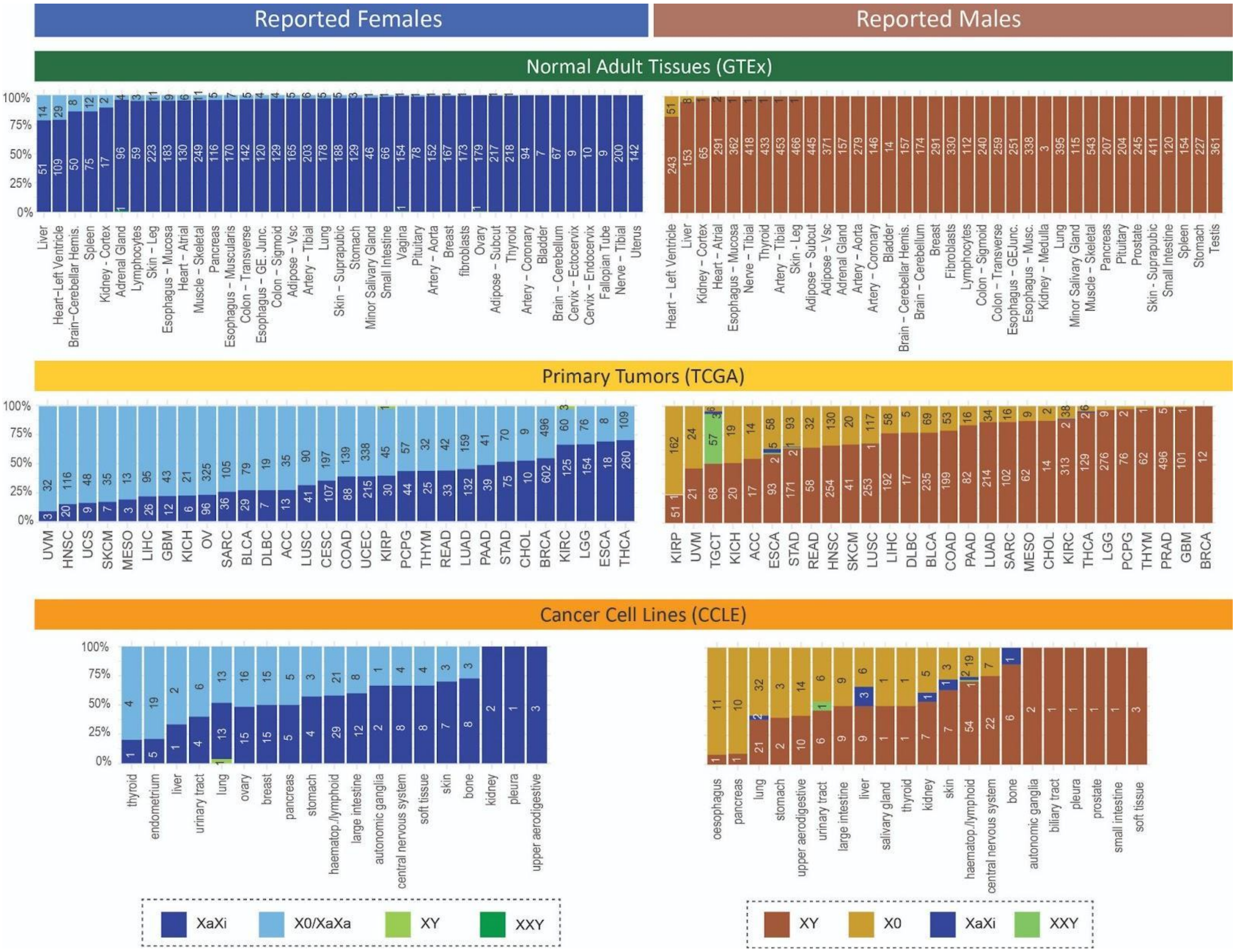

**Fig. S4. Sex Chromosome Complement in primary tumors versus adjacent normal tissues**. Violin plots showing the difference between expression of sex chromosome complement inference genes in primary tumor tissue and the adjacent normal tissue from the same patients. **(A)** XIST expression shown in reported female patients across tumor types. **(B)** DDX3Y expression shown in reported male patients. **(C)** Difference between expression in primary tumor tissue and adjacent normal tissue of XIST in female patients and DDX3Y in male patients. Blue dots indicate patients for which XIST or DDX3Y were not expressed in adjacent normal and expressed in the primary tumor, green dots for which they were expressed in adjacent normal but lost in the primary tumor, and light blue and pink dots show patients that did not change in their XIST or DDX3Y status between adjacent normal and primary tumor tissues.

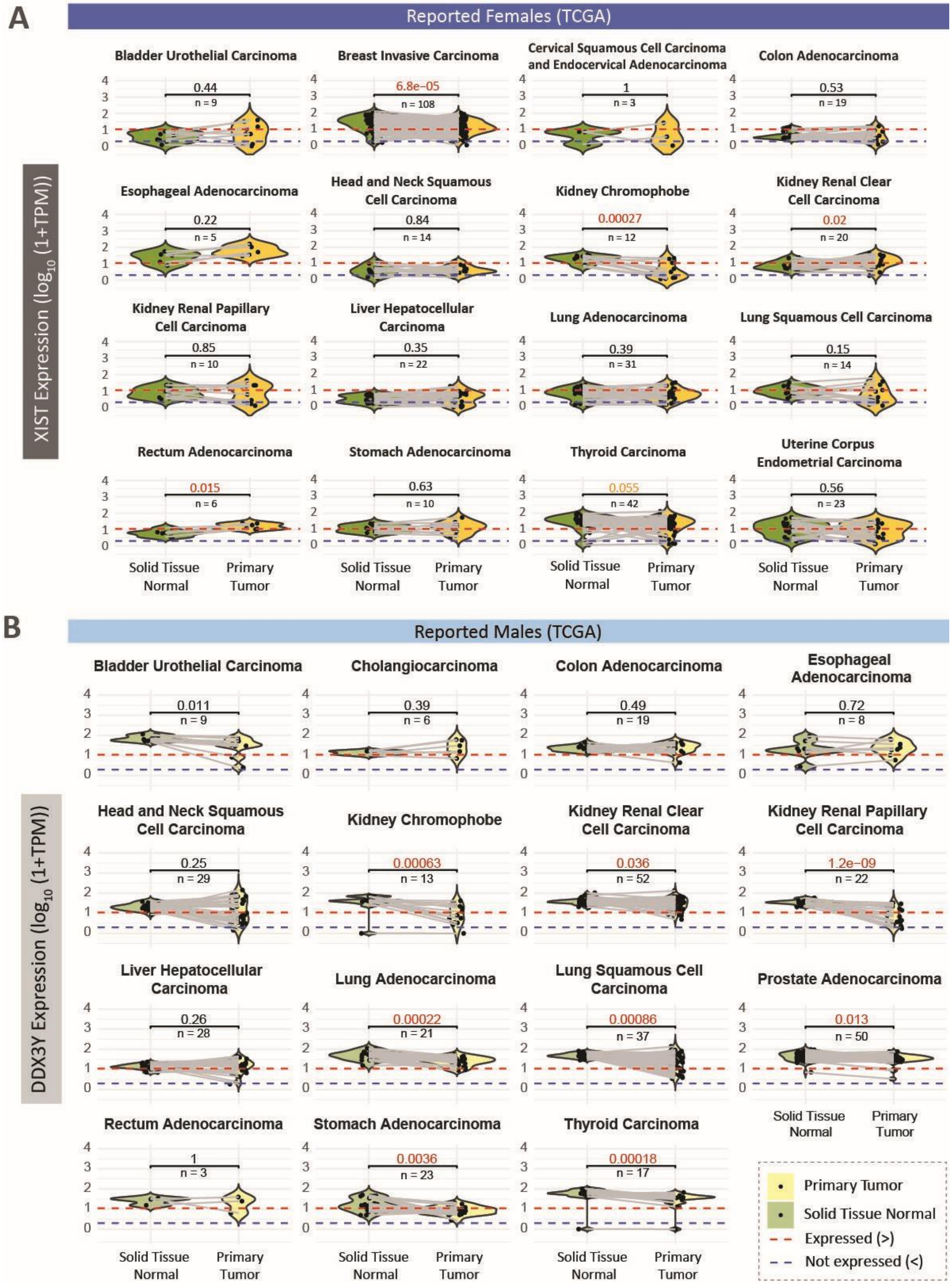
A
Reported Females (TCGA)
XIST Expression (log10 (1+TPM))
Bladder Urothelial Carcinoma
0.44
n = 9
Breast Invasive Carcinoma
6.8e−05
n = 108
Cervical Squamous Cell Carcinoma and Endocervical Adenocarcinoma
1
n = 3
Colon Adenocarcinoma
0.53
n = 19
Esophageal Adenocarcinoma
0.22
n = 5
Head and Neck Squamous Cell Carcinoma
0.84
n = 14
Kidney Chromophobe
0.00027
n = 12
Kidney Renal Clear Cell Carcinoma
0.02
n = 20
Kidney Renal Papillary Cell Carcinoma
0.85
n = 10
Liver Hepatocellular Carcinoma
0.35
n = 22
Lung Adenocarcinoma
0.39
n = 31
Lung Squamous Cell Carcinoma
0.15
n = 14
Rectum Adenocarcinoma
0.015
n = 6
Stomach Adenocarcinoma
0.63
n = 10
Thyroid Carcinoma
0.055
n = 42
Uterine Corpus Endometrial Carcinoma
0.56
n = 23
Solid Tissue Normal
Primary Tumor
B
Reported Males (TCGA)
DDX3Y Expression (log10 (1+TPM))
Bladder Urothelial Carcinoma
0.011
n = 9
Cholangiocarcinoma
0.39
n = 6
Colon Adenocarcinoma
0.49
n = 19
Esophageal Adenocarcinoma
0.72
n = 8
Head and Neck Squamous Cell Carcinoma
0.25
n = 29
Kidney Chromophobe
0.00063
n = 13
Kidney Renal Clear Cell Carcinoma
0.036
n = 52
Kidney Renal Papillary Cell Carcinoma
1.2e−09
n = 22
Liver Hepatocellular Carcinoma
0.26
n = 28
Lung Adenocarcinoma
0.00022
n = 21
Lung Squamous Cell Carcinoma
0.00086
n = 37
Prostate Adenocarcinoma
0.013
n = 50
Rectum Adenocarcinoma
1
n = 3
Stomach Adenocarcinoma
0.0036
n = 23
Thyroid Carcinoma
0.00018
n = 17
Solid Tissue Normal
Primary Tumor
Primary Tumor
Solid Tissue Normal
Expressed (>)
Not expressed (<)

C

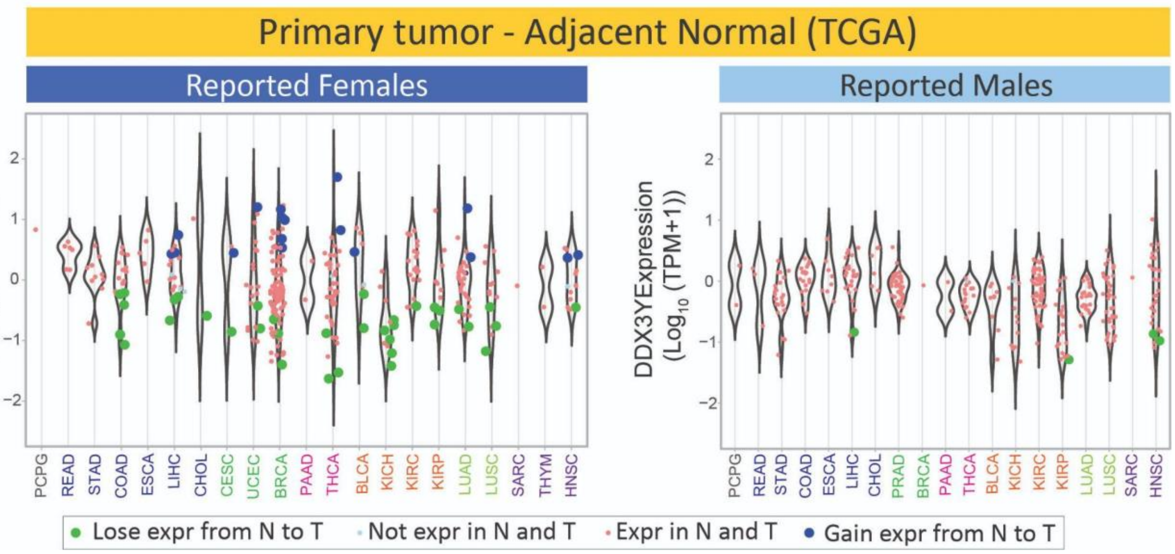
Primary tumor - Adjacent Normal (TCGA)
Reported Females
Reported Males
XIST Expression (Log10 (TPM+1))
DDX3YExpression (Log10 (TPM+1))
PCPG READ STAD COAD ESCA LIHC CHOL CESC UCEC BRCA PAAD THCA BLCA KICH KIRC KIRP LUAD LUSC SARC THYM HNSC
PCPG READ STAD COAD ESCA LIHC CHOL PRAD BRCA PAAD THCA BLCA KICH KIRC KIRP LUAD LUSC SARC HNSC
Lose expr from N to T
Not expr in N and T
Expr in N and T
Gain expr from N to T

**Fig. S5.** ***XIST*** **and** ***DDX3Y*** **gene expression and relative protein abundance in normal tissues, primary tumor tissues, and cancer cell lines in all tissues. (A)** Violin plots showing the distribution of XIST expression in reported males versus reported females in GTEx (normal non-cancerous tissues), TCGA (primary tumor tissues), and CCLE (cancer cell lines) data sets. Red dashed line, minimum threshold for expression (≥ 10 transcripts per million), blue dashed line, maximum threshold for no expression (≤ 1 transcript per million), between dashed lines, intermediate expression. **(B)** Violin plots showing the distribution of DDX3Y in both sexes in three datasets in gene expression and relative protein abundance. **(C)** Pie chart showing the proportion of inferred sex chromosome complements within reported males and reported females.

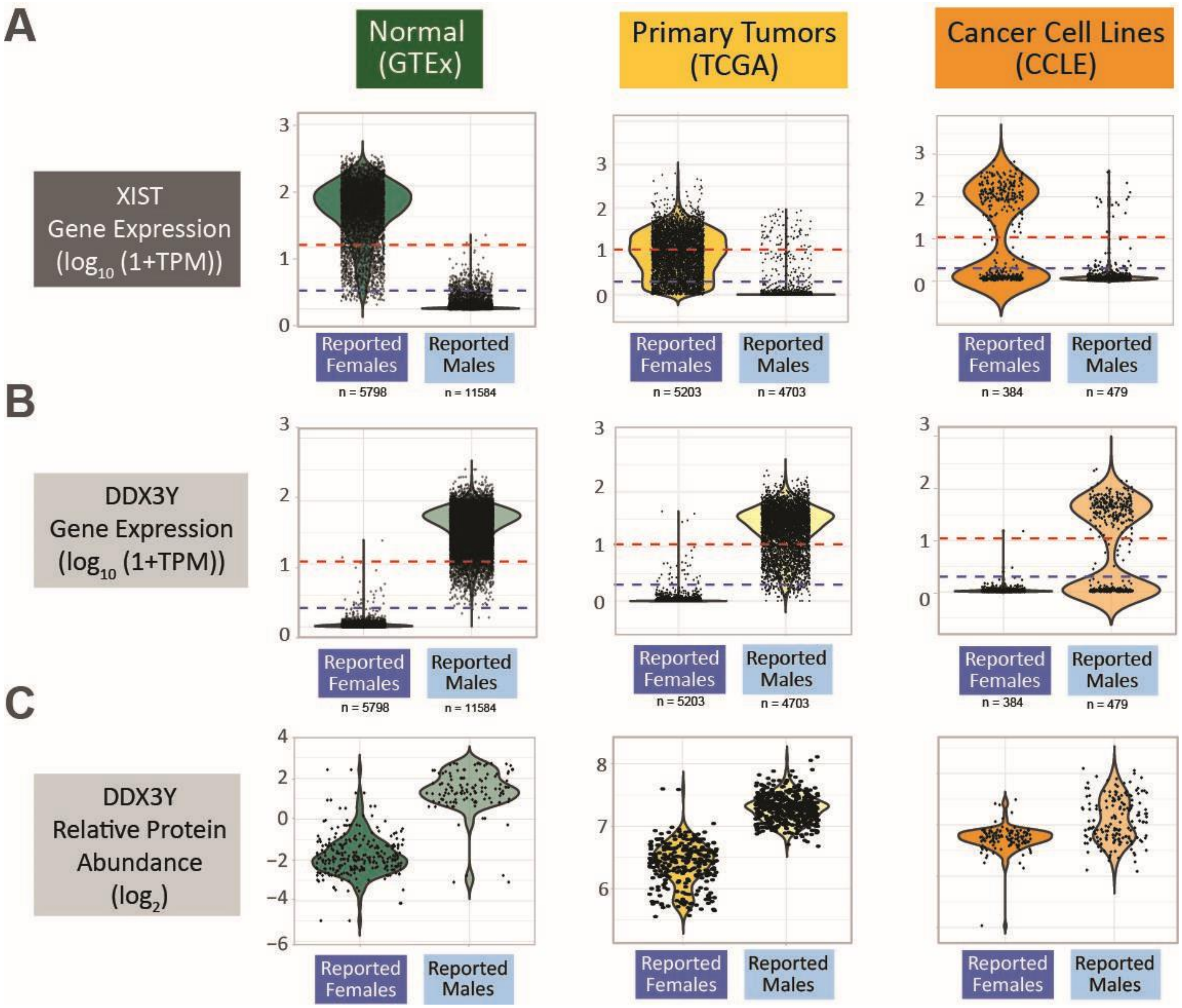

**Fig. S6. High correlation between gene expression and protein expression**. Chromosome Y genes that were detected in mass spectrometry proteomics (DDX3Y, EIF1AY, and RPS4Y1) are plotted against gene expression data for those genes. Correlation coefficients are shown in the top left corner, sex and cancer status are marked using point color and shape.

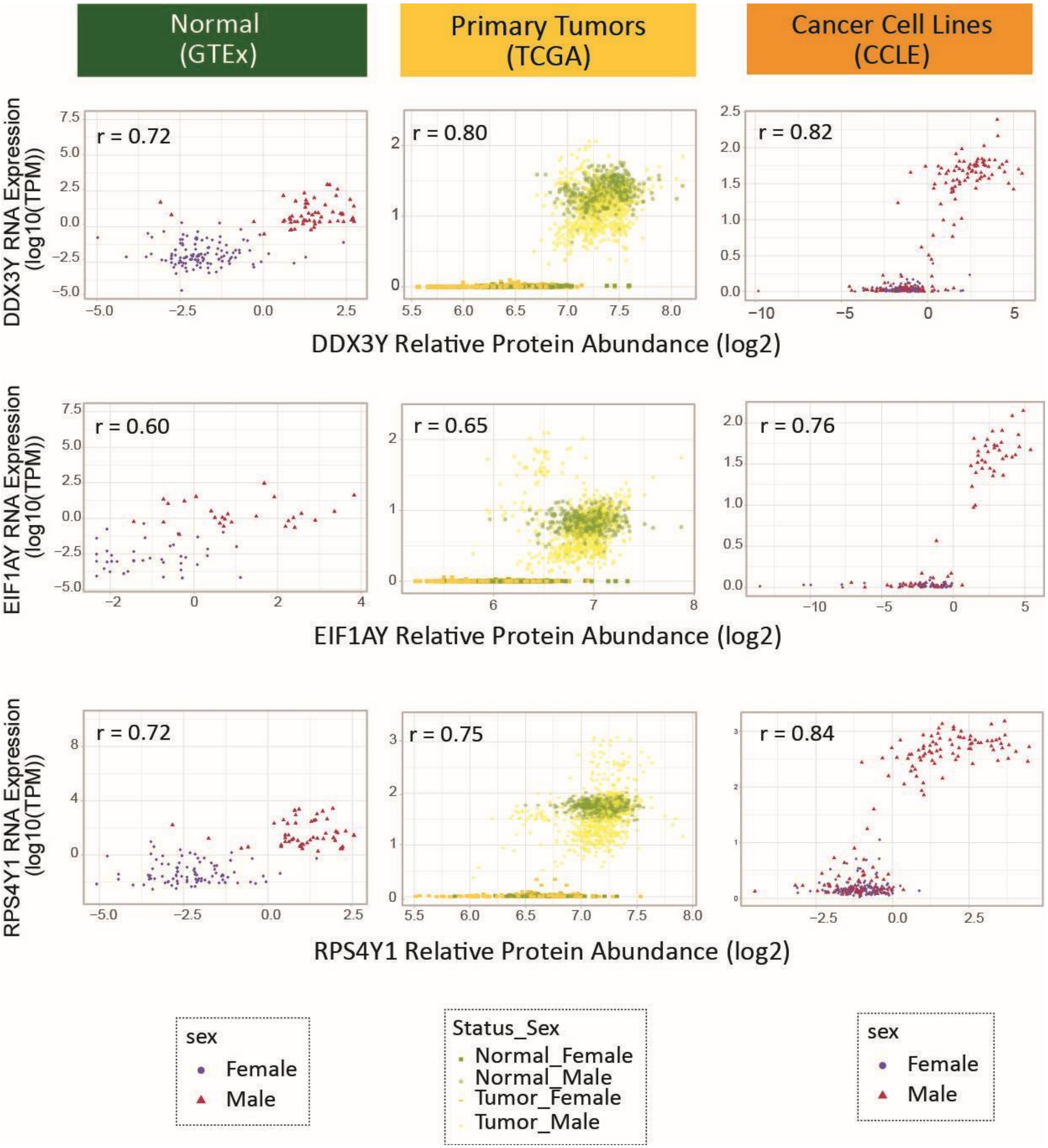

**Fig. S7. Age differences between sample groups based on their XIST or chromosome Y expression levels.** **(A)** Linear modeling prediction of sex chromosome gene expression with age as a dependent variable. **(B)** Violin plots showing the distribution of age of the individuals from which samples were taken. Mean age and statistical differences between groups are labelled.

**A**

| Dataset | Reported Sex | Expression | Estimate | Std Err | t | Pr(> \|t\|) | Significance |
|---|---|---|---|---|---|---|---|
| GTEX | Female | *XIST* TPM | -0.22 | 0.05 | -4.26 | <2e-16 | *** |
| GTEX | Male | chrY gene avg TPM | -0.16 | 0.01 | -10.61 | <2e-16 | *** |
| TCGA | Female | *XIST* TPM | -0.20 | 0.03 | -6.50 | 8.8E-11 | *** |
| TCGA | Male | chrY gene avg TPM | -0.31 | 0.04 | -7.26 | 4.3E-13 | *** |
| CCLE | Female | *XIST* TPM | -0.82 | 0.32 | -2.53 | 0.01 | * |
| CCLE | Male | chrY gene avg TPM | -0.71 | 0.14 | -4.77 | 2.6E-06 | * |

**B**

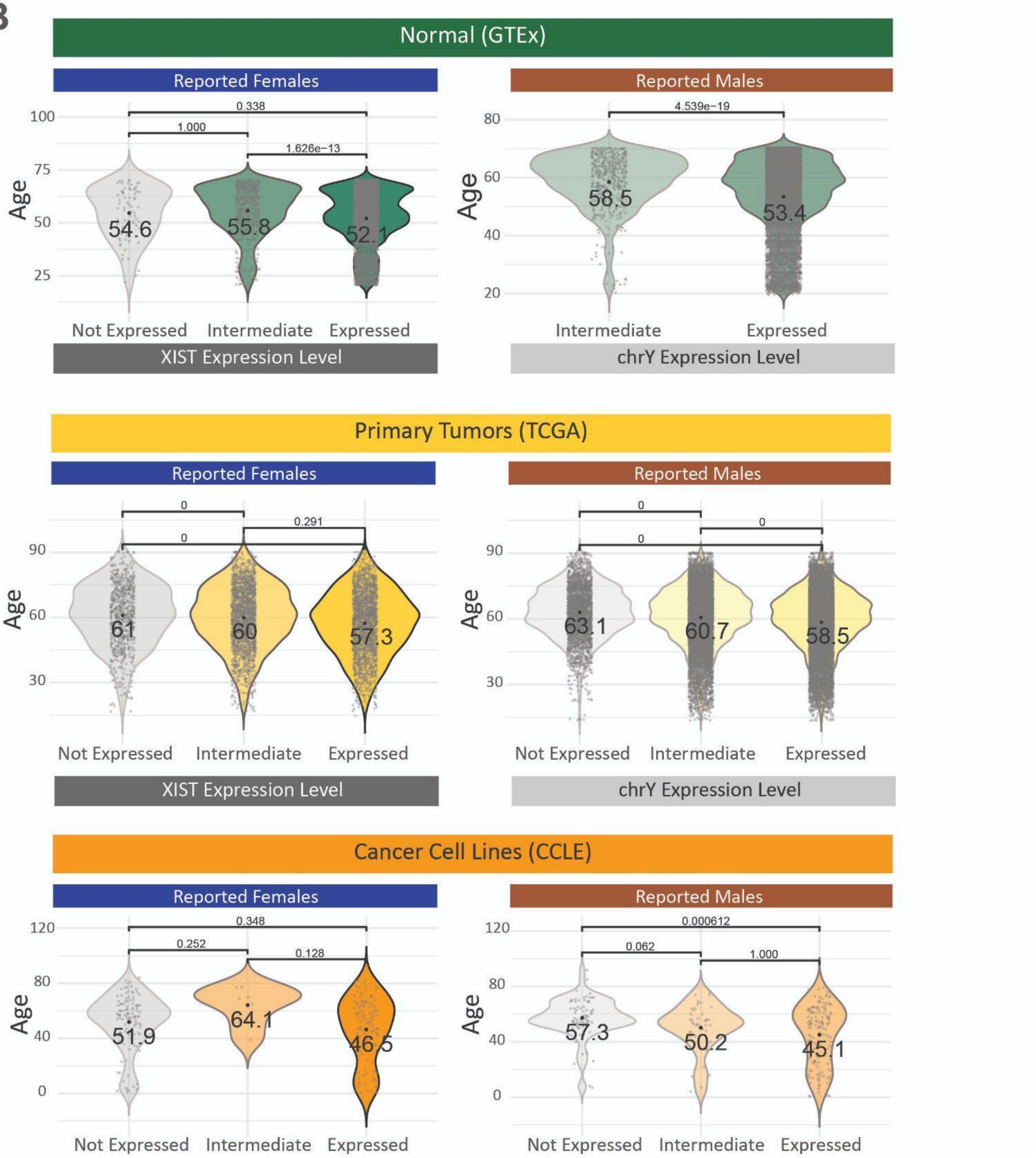

**Fig. S8. Correlation of sex chromosome based gene expression profiles across tissues.** For tissues that had at least three tumor samples with each sex chromosome complement being compared, log-transformed fold changes (ratios of average expression) were clustered and showed no distinct subclusters within the genes. Correlation coefficients were calculated between the log fold change profiles comparing the tissues pairwise showing only positive correlations across the tissues.

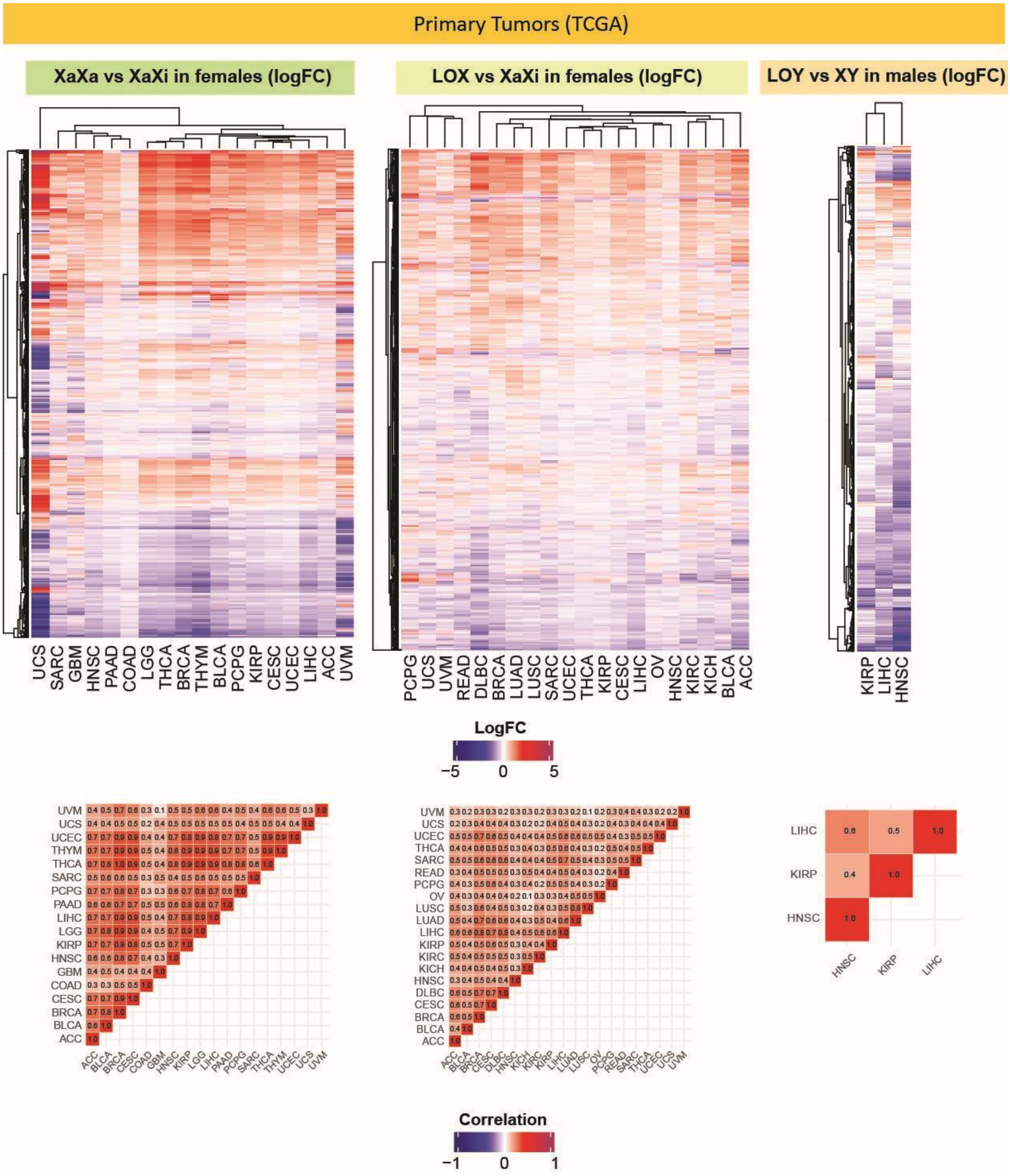

**Fig. S9. Genes affected by loss of sex chromosomes consistently across normal adult tissue types.** Consistent gene expression changes that were in the same direction and statistically significant differential expression (p-value < 0.05) in at least 60% of tissues in GTEx. Genes changing: (**A**) with loss of *XIST* expression indicating loss of the X chromosome or loss of X chromosome inactivation(X0/XaXa : XaXi) in tissues from female patients, (**B**) with loss of the Y chromosome (X0 : XY) in tissues from male patients. Genes are annotated with known functions and associations with hallmarks of cancer.

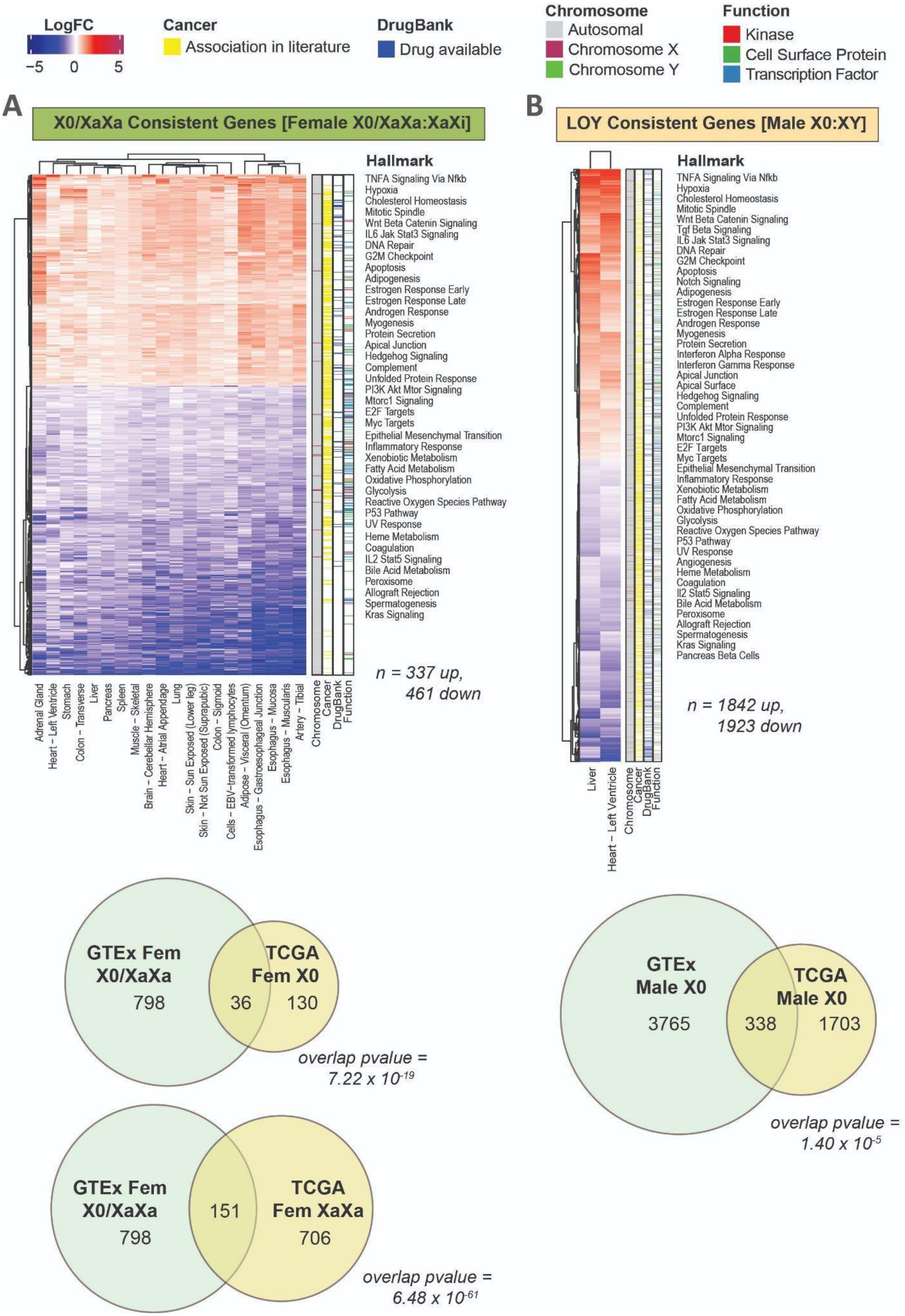
LogFC
−5 0 5
Cancer
Association in literature
DrugBank
Drug available
Chromosome
Autosomal
Chromosome X
Chromosome Y
Function
Kinase
Cell Surface Protein
Transcription Factor
A
X0/XaXa Consistent Genes [Female X0/XaXa:XaXi]
Hallmark
TNFA Signaling Via Nfkb
Hypoxia
Cholesterol Homeostasis
Mitotic Spindle
Wnt Beta Catenin Signaling
IL6 Jak Stat3 Signaling
DNA Repair
G2M Checkpoint
Apoptosis
Adipogenesis
Estrogen Response Early
Estrogen Response Late
Androgen Response
Myogenesis
Protein Secretion
Apical Junction
Hedgehog Signaling
Complement
Unfolded Protein Response
PI3K Akt Mtor Signaling
Mtorc1 Signaling
E2F Targets
Myc Targets
Epithelial Mesenchymal Transition
Inflammatory Response
Xenobiotic Metabolism
Fatty Acid Metabolism
Oxidative Phosphorylation
Glycolysis
Reactive Oxygen Species Pathway
P53 Pathway
UV Response
Heme Metabolism
Coagulation
IL2 Stat5 Signaling
Bile Acid Metabolism
Peroxisome
Allograft Rejection
Spermatogenesis
Kras Signaling
n = 337 up,
461 down
Adrenal Gland
Heart – Left Ventricle
Stomach
Colon – Transverse
Liver
Pancreas
Spleen
Muscle – Skeletal
Brain – Cerebellar Hemisphere
Heart – Atrial Appendage
Lung
Skin – Sun Exposed (Lower leg)
Skin – Not Sun Exposed (Suprapubic)
Colon – Sigmoid
Cells – EBV-transformed lymphocytes
Adipose – Visceral (Omentum)
Esophagus – Gastroesophageal Junction
Esophagus – Mucosa
Esophagus – Muscularis
Artery – Tibial
Chromosome
Cancer
DrugBank
Function
B
LOY Consistent Genes [Male X0:XY]
Hallmark
TNFA Signaling Via Nfkb
Hypoxia
Cholesterol Homeostasis
Mitotic Spindle
Wnt Beta Catenin Signaling
Tgf Beta Signaling
IL6 Jak Stat3 Signaling
DNA Repair
G2M Checkpoint
Apoptosis
Notch Signaling
Adipogenesis
Estrogen Response Early
Estrogen Response Late
Androgen Response
Myogenesis
Protein Secretion
Interferon Alpha Response
Interferon Gamma Response
Apical Junction
Apical Surface
Hedgehog Signaling
Complement
Unfolded Protein Response
PI3K Akt Mtor Signaling
Mtorc1 Signaling
E2F Targets
Myc Targets
Epithelial Mesenchymal Transition
Inflammatory Response
Xenobiotic Metabolism
Fatty Acid Metabolism
Oxidative Phosphorylation
Glycolysis
Reactive Oxygen Species Pathway
P53 Pathway
UV Response
Angiogenesis
Heme Metabolism
Coagulation
Il2 Stat5 Signaling
Bile Acid Metabolism
Peroxisome
Allograft Rejection
Spermatogenesis
Kras Signaling
Pancreas Beta Cells
n = 1842 up,
1923 down
Liver
Heart – Left Ventricle
Chromosome
Cancer
DrugBank
Function
GTEx Fem X0/XaXa
798
36
TCGA Fem X0
130
overlap pvalue = 7.22 x 10-19
GTEx Fem X0/XaXa
798
151
TCGA Fem XaXa
706
overlap pvalue = 6.48 x 10-61
GTEx Male X0
3765
338
TCGA Male X0
1703
overlap pvalue = 1.40 x 10-5

**Fig. S10. Genes affected by loss of sex chromosomes consistently across cancer cell lines from different tumor tissue types.** Consistent gene expression changes that were in the same direction and statistically significant differential expression (p-value < 0.05) in at least 60% of tissues in CCLE. Upset plot showing the number of differentially expressed genes in each tissue type. Genes changing: (**A, B**) with loss of *XIST* expression indicating loss of the X chromosome or loss of X chromosome inactivation (X0/XaXa : XaXi) in tissues from female patients, (**C, D**) with loss of the Y chromosome (X0 : XY) in tissues from male patients.

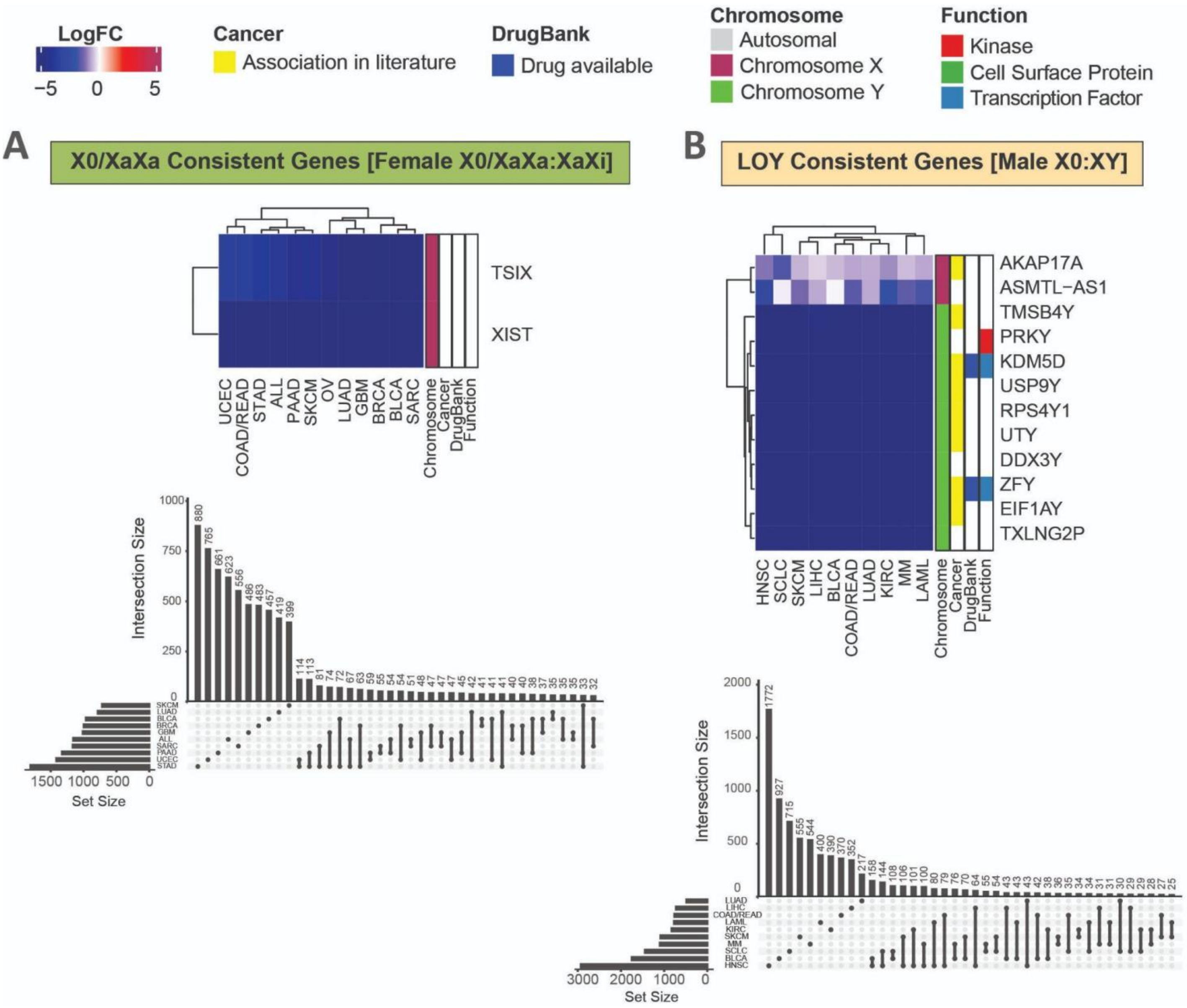

**Fig. S11. ANOVA/PLSDA analysis at different thresholds.** ANOVA analysis of genes differentially expressed between two or more sample groups determined from sex chromosome gene expression and DNA ploidy of the sex chromosomes was conducted, the top most significant genes are mapped according to those genes' expression using partial least squares discriminant analysis. Analysis was conducted in normal adult tissues (GTEx), primary tumor tissues (TCGA), and cancer cell lines (CCLE).

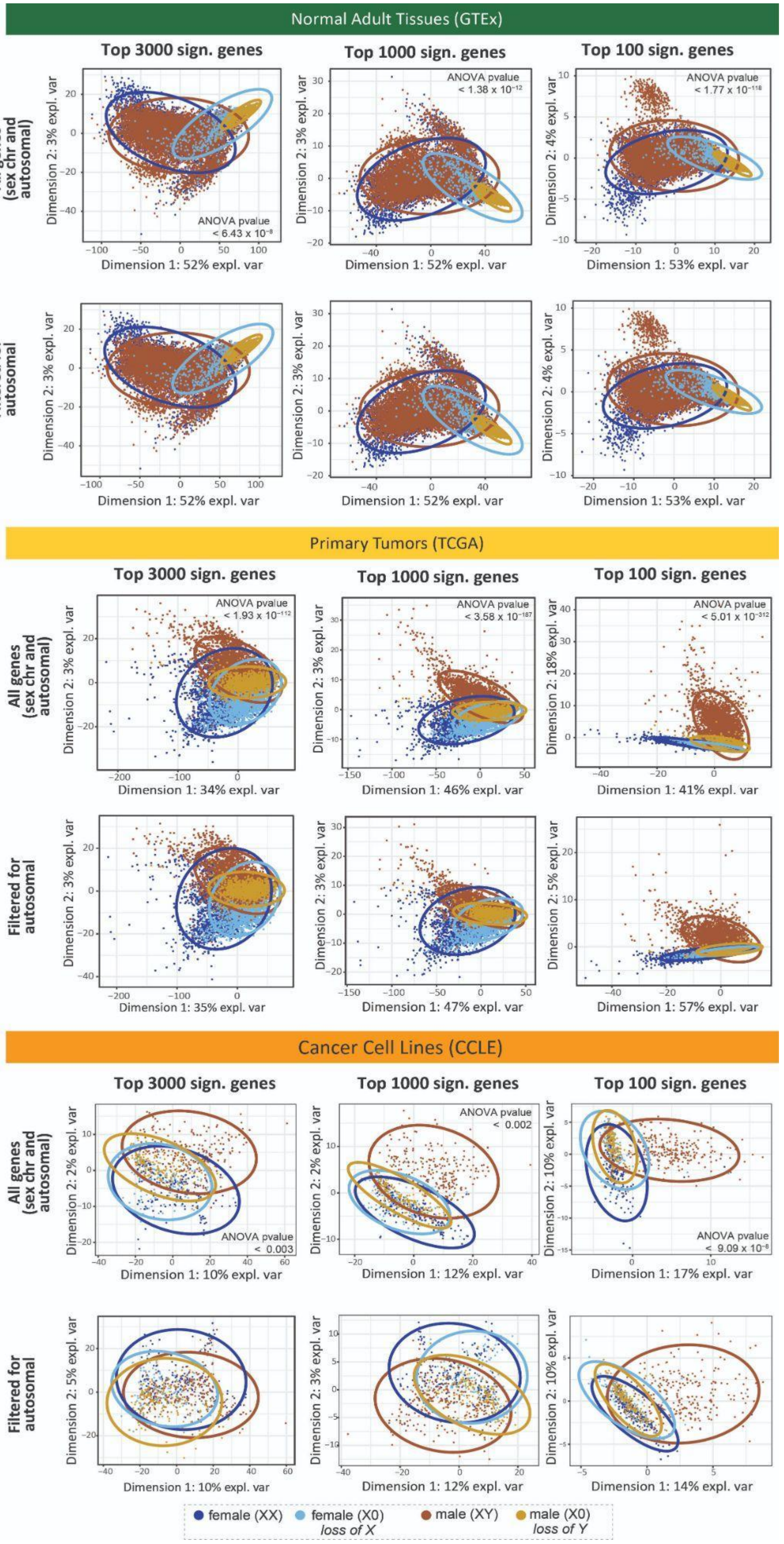

Normal Adult Tissues (GTEx)
Top 3000 sign. genes
Top 1000 sign. genes
Top 100 sign. genes
All genes (sex chr and autosomal)
Filtered for autosomal
ANOVA pvalue < 6.43 x $10^{-8}$
ANOVA pvalue < 1.38 x $10^{-12}$
ANOVA pvalue < 1.77 x $10^{-118}$
Dimension 1: 52% expl. var
Dimension 1: 53% expl. var
Dimension 2: 3% expl. var
Dimension 2: 4% expl. var
Primary Tumors (TCGA)
Top 3000 sign. genes
Top 1000 sign. genes
Top 100 sign. genes
All genes (sex chr and autosomal)
Filtered for autosomal
ANOVA pvalue < 1.93 x $10^{-112}$
ANOVA pvalue < 3.58 x $10^{-187}$
ANOVA pvalue < 5.01 x $10^{-312}$
Dimension 1: 34% expl. var
Dimension 1: 46% expl. var
Dimension 1: 41% expl. var
Dimension 1: 35% expl. var
Dimension 1: 47% expl. var
Dimension 1: 57% expl. var
Dimension 2: 3% expl. var
Dimension 2: 18% expl. var
Dimension 2: 5% expl. var
Cancer Cell Lines (CCLE)
Top 3000 sign. genes
Top 1000 sign. genes
Top 100 sign. genes
All genes (sex chr and autosomal)
Filtered for autosomal
ANOVA pvalue < 0.003
ANOVA pvalue < 0.002
ANOVA pvalue < 9.09 x $10^{-8}$
Dimension 1: 10% expl. var
Dimension 1: 12% expl. var
Dimension 1: 17% expl. var
Dimension 1: 14% expl. var
Dimension 2: 2% expl. var
Dimension 2: 10% expl. var
Dimension 2: 5% expl. var
Dimension 2: 3% expl. var
female (XX)
female (X0) loss of X
male (XY)
male (X0) loss of Y

**Fig. S12. Aneuploidy in tumors based on sex chromosome changes**. **(A)** Aneuploidy score across SCC groups. **(B)** Expression of *APC* gene in SCC groups of tumors. Comparisons that are statistically significant differential expression are labelled (adjusted p-value with age as a covariate < 0.05).

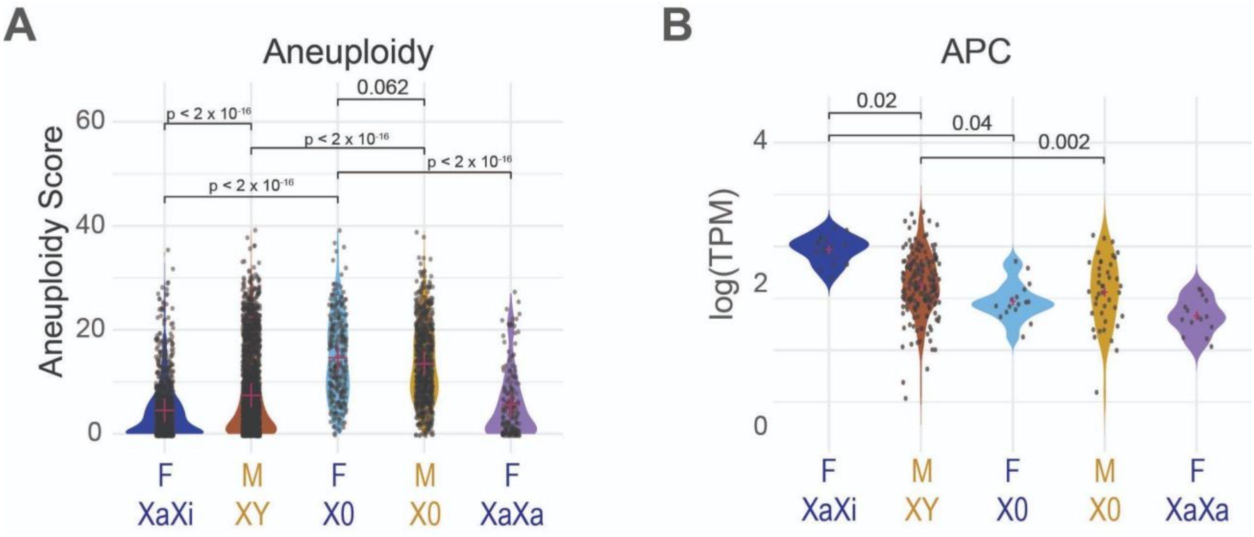


**Fig. S13. Venn diagrams comparing differential gene expression between profiles of sex chromosome changes**. Overlap between genes differentially expressed (adjusted p-value < 0.05) between tumors from females that have reactivated one X chromosome compared to the expected genotype (XaXa : XaXi) , labelled 'XaXa', genes differentially expressed between tumors from females that have lost an X chromosome (XaXi : X0), labelled 'LOX', and genes differentially expressed between tumor samples from males who have lost a Y chromosome compared to the expected genotype (XY : X0), labelled 'LOY'.

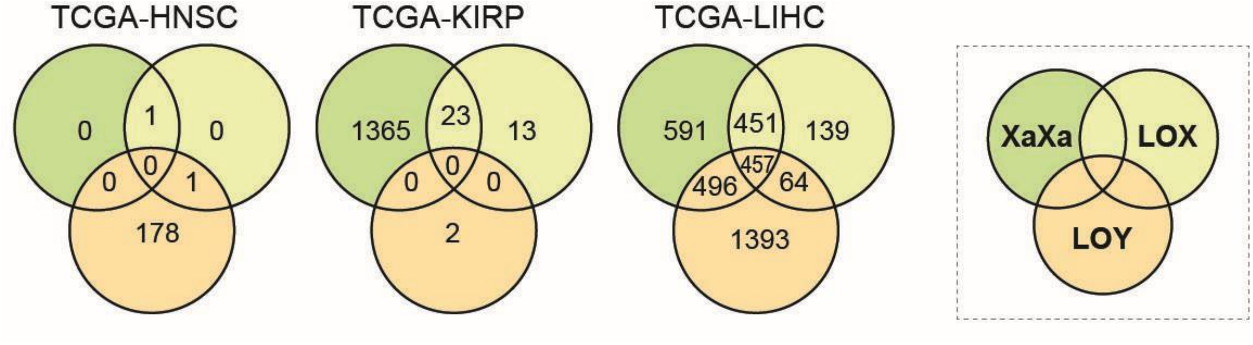

**Table S1. (A)** Survival analysis for patients from TCGA based on their sex and sex chromosome complement. **(B)** Pairwise comparison of survival curves.

**Survival properties**

| | Records | Events | Mean (d,y) | SE [Mean] (d) | Median (d,y) | Cox PH | Cox PH p-value |
|---|---|---|---|---|---|---|---|
| **XaXi (fem)** | 1200 | 219 | 4641.4 (12.7) | 271.7 | 4445 (12.2) | ref | ref |
| **XaXa (fem)** | 208 | 44 | 6638.6 (18.2) | 648.1 | not reached | 1.16 | 0.38 |
| **XY (male)** | 2664 | 726 | 4134.9 (11.3) | 269.4 | 3200 (8.8) | 1.57 | $4.5 \times 10^{-9}$ *** |
| **X0 (fem)** | 264 | 100 | 2553.9 (7.0) | 171.5 | 1691 (4.6) | 2.04 | $3.0 \times 10^{-9}$ *** |
| **X0 (male)** | 885 | 321 | 3400.1 (9.3) | 329.3 | 2043 (5.6) | 2.18 | $< 2 \times 10^{-16}$ *** |

**Pairwise differences in survival (p-values)**

| | X0 (fem) | XaXa (fem) | XaXi (fem) | X0 (male) |
|---|---|---|---|---|
| **XaXa (fem)** | 0.0025 | - | - | - |
| **XaXi (fem)** | $8.8 \times 10^{-9}$ | 0.44 | - | - |
| **X0 (male)** | 0.58 | 0.00012 | $< 2 \times 10^{-16}$ | - |
| **XY (male)** | 0.02 | 0.061 | $1.2 \times 10^{-8}$ | $2.1 \times 10^{-6}$ |
